# Human neutralizing antibodies against SARS-CoV-2 require intact Fc effector functions and monocytes for optimal therapeutic protection

**DOI:** 10.1101/2020.12.28.424554

**Authors:** Emma S. Winkler, Pavlo Gilchuk, Jinsheng Yu, Adam L. Bailey, Rita E. Chen, Seth J. Zost, Hyesun Jang, Ying Huang, James D. Allen, James Brett Case, Rachel E. Sutton, Robert H. Carnahan, Tamarand L. Darling, Adrianus C. M. Boon, Matthias Mack, Richard D. Head, Ted M. Ross, James E. Crowe, Michael S. Diamond

## Abstract

SARS-CoV-2 has caused the global COVID-19 pandemic. Although passively delivered neutralizing antibodies against SARS-CoV-2 show promise in clinical trials, their mechanism of action *in vivo* is incompletely understood. Here, we define correlates of protection of neutralizing human monoclonal antibodies (mAbs) in SARS-CoV-2-infected animals. Whereas Fc effector functions are dispensable when representative neutralizing mAbs are administered as prophylaxis, they are required for optimal protection as therapy. When given after infection, intact mAbs reduce SARS-CoV-2 burden and lung disease in mice and hamsters better than loss-of-function Fc variant mAbs. Fc engagement of neutralizing antibodies mitigates inflammation and improves respiratory mechanics, and transcriptional profiling suggests these phenotypes are associated with diminished innate immune signaling and preserved tissue repair. Immune cell depletions establish that neutralizing mAbs require monocytes for therapeutic efficacy. Thus, potently neutralizing mAbs require Fc effector functions for maximal therapeutic benefit during therapy to modulate protective immune responses and mitigate lung disease.

## INTRODUCTION

Severe Acute Respiratory Syndrome Coronavirus-2 (SARS-CoV-2) is the recently emerged RNA virus responsible for the Coronavirus Disease 2019 (COVID-19) pandemic that has led to over 52 million infections and over 1.29 million deaths (Dong et al., 2020). Clinical disease is variable, ranging from asymptomatic infection to multi-organ failure and death. The need for effective countermeasures of COVID-19 is urgent, as only dexamethasone and remdesivir have demonstrated clinical efficacy in specific indications. One promising approach for both the prevention and treatment COVID-19 has been the development of SARS-CoV-2-neutralizing monoclonal antibodies (mAbs) isolated from the B cells of individuals with recent SARS-CoV-2 infection (Abraham, 2020; Marovich et al., 2020). Multiple neutralizing mAbs directed against the spike (S) glycoprotein of SARS-CoV-2 have been identified that target non-overlapping epitopes and show differences in neutralization potency (Alsoussi et al., 2020; Cao et al., 2020; Hansen et al., 2020; Ju et al., 2020; Liu et al., 2020; Noy-Porat et al., 2020; Pinto et al., 2020; Robbiani et al., 2020; Zost et al., 2020a). The majority of these mAbs bind to the S1 subunit, specifically the receptor binding domain (RBD), and inhibit virus attachment to its principal cell surface receptor, angiotensin converting enzyme (ACE)2.

Prophylactic and therapeutic efficacy of anti-S mAbs has been demonstrated *in vivo* in murine, hamster, and non-human primate models of SARS-CoV-2 pathogenesis (Alsoussi et al., 2020; Baum et al., 2020; Fagre et al., 2020; Hansen et al., 2020; Hassan et al., 2020; Kreye et al., 2020; Rogers et al., 2020; Shi et al., 2020; Zost et al., 2020a), with varying degrees of reduction in viral burden and dampened inflammation of the lung. However, the mechanisms of protection *in vivo* can be due to multiple factors including direct virus neutralization and engagement of complement or Fc gamma receptors (FcγRs) on leukocytes. Fc effector functions of antibodies can promote immune-mediated clearance mechanisms (by antibody-dependent cellular cytotoxicity, antibody-dependent cellular phagocytosis, and complement activation), enhance antigen presentation and CD8^+^ T cell responses, and reshape inflammation through engagement of individual FcγRs on specific hematopoietic cells (Lu et al., 2018). In contrast, under certain circumstances, Fc-FcγR interactions can promote antibody-dependent enhancement of virus infection (ADE) (Halstead, 1994) or pathological immune skewing (Bolles et al., 2011; Ruckwardt et al., 2019), which is at least a theoretical concern of antibody-based therapies and vaccines against SARS-CoV-2 (Diamond and Pierson, 2020). Thus, a more thorough understanding of the contribution of Fc effector functions in the context of antibody-based therapies is needed.

Here, we use the K18-hACE2 transgenic mouse model of SARS-CoV-2 pathogenesis (Golden et al., 2020; Winkler et al., 2020) and a Fc region genetic variant form of IgG (LALA-PG) of a potent RBD-binding neutralizing mAb (COV2-2050) that cannot engage FcγRs or complement to define the role of Fc effector functions in antibody protection. We find that Fc effector functions are dispensable when neutralizing mAbs are administered as prophylaxis, but unexpectedly, they are required for optimal protection when given as post-exposure therapy. When administered after SARS-CoV-2 infection, intact but not LALA-PG mAbs reduce viral burden and lung disease. Fc engagement by antibodies decreases immune cell infiltration and levels of inflammatory cytokines, and this activity is linked to improved respiratory mechanics and outcome. RNA sequencing analysis of lung homogenates reveals distinct gene signatures associated with Fc engagement including decreased innate immune signaling and extracellular matrix remodeling but maintained expression of genes that mediate tissue repair. Immune cell depletions showed that neutralizing mAbs require monocytes for therapeutic efficacy. We confirmed these findings in mice with additional RBD-binding neutralizing mAbs (COV2-3025, COV2-2072, and COV2-2381) as well as in hamsters, a second mammalian species, as Fc effector functions of a neutralizing mAb also are required to prevent weight loss, control viral infection, and limit inflammation. Overall, these studies establish that Fc effector functions of neutralizing antibody are necessary for an optimal therapeutic outcome after SARS-CoV-2 infection.

## RESULTS

### Pre-exposure protection against SARS-CoV-2 infection in mice by a neutralizing mAb does not require Fc effector functions

Passive transfer of neutralizing mAbs targeting the S protein confers protection in multiple pre-clinical models of SARS CoV-2 infection (Alsoussi et al., 2020; Baum et al., 2020; Hansen et al., 2020; Hassan et al., 2020; Kreye et al., 2020; Rogers et al., 2020; Zost et al., 2020a). However, *in vivo* antiviral efficacy can be due to multiple mechanisms including Fab-dependent direct virus neutralization and Fc-dependent engagement that promotes opsonization of virus, clearance of virally-infected cells, and modulation of innate and adaptive immune responses (Bournazos et al., 2019; Bournazos et al., 2014b; DiLillo et al., 2016; DiLillo et al., 2014; Fox et al., 2019). To evaluate the contribution of Fc effector functions, we introduced loss-of-function LALA-PG (L234A, L235A, and P329G) mutations into the human IgG1 heavy chain of COV2-2050, a previously described neutralizing human mAb that binds the RBD of S and blocks ACE2 binding (Zost et al., 2020a), to abolish antibody Fc interactions with FcγRs and complement proteins (Lo et al., 2017). Intact or LALA-PG variants of COV2-2050 neutralized SARS-CoV-2 equivalently in cell culture (**Fig 1A**), and introduction of the LALA-PG mutation prevented binding to mouse FcγRI and FcγRIV (**Fig 1B**). To test their relative efficacy *in vivo*, we used the K18-hACE2 transgenic mouse model of SARS-CoV-2 pathogenesis in which human ACE2 expression is driven by the cytokeratin-18 gene promoter (McCray et al., 2007; Winkler et al., 2020). In prophylaxis studies, K18-hACE2 transgenic mice received decreasing doses of COV2-2050 or COV2-2050 LALA-PG (200 μg (10 mg/kg), 40 μg (2 mg/kg), 8 μg (0.4 mg/kg), or 1.6 μg (0.08 mg/kg)) by intraperitoneal injection 16 h prior to intranasal inoculation with SARS-CoV-2 (10^3^ PFU, strain 2019 n-CoV/USA_WA1/2020). An isotype control human mAb (DENV-2D22) was delivered at a single 200 μg dose for comparison. At the time of SARS-CoV-2 challenge, the serum levels of COV2-2050 or COV2-2050 LALA-PG were equivalent (**Fig 1C**). As expected, lower doses of COV2-2050 resulted in decreasing neutralizing antibody titers in serum at the time of virus inoculation (**Fig 1D**).

**Figure 1.**
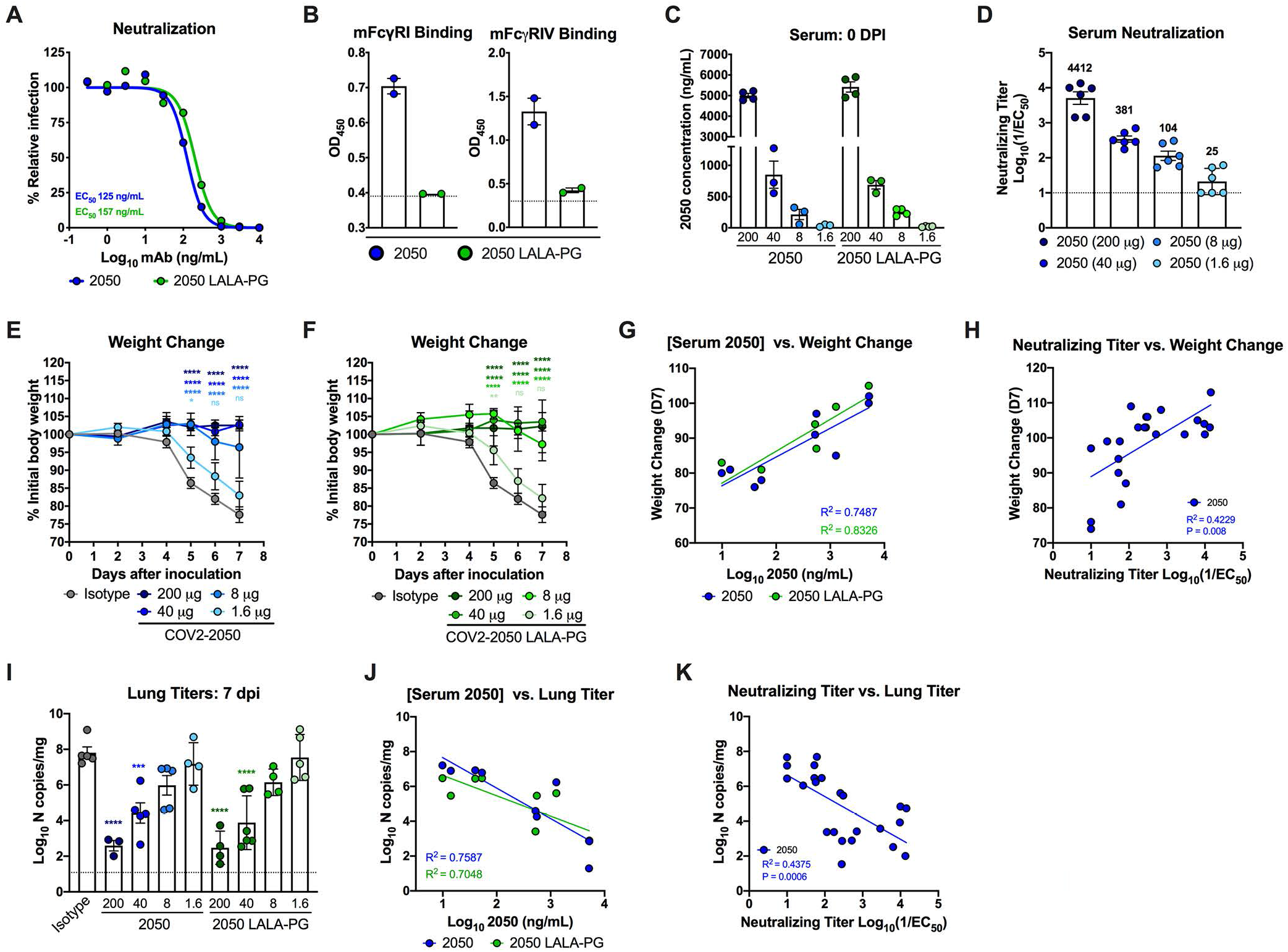
Neutralizing activity is sufficient for prophylactic efficacy of mAbs against SARS-CoV-2 infection in K18-hACE2 transgenic mice. (**A**) Anti-SARS-CoV-2 mAbs (COV2-2050 and COV2-2050 LALA-PG) were incubated with 10^2^ focus-forming units (FFU) of SARS-CoV-2 for 1 h at 37°C followed by addition of mAb-virus mixture to Vero E6 cells. Wells containing mAb were compared to wells without mAb to determine relative infection. One experiment of two, with similar results, is shown. The mean of two technical replicates is shown. (**B**) Binding of COV2-2050 or COV2-2050 LALA-PG to recombinant mouse FcγRI and FcγRIV as measured by ELISA (two independent experiments). For **C-J,** eight-week-old female and male K18-hACE2 transgenic mice received 200 μg, 40 μg, 8 μg, or 1.6 μg of COV2-2050 or COV2-2050 LALA-PG one day prior to intranasal inoculation with 10^3^ PFU of SARS-CoV-2. (**C**) Serum concentrations (ng/mL) of COV2-2050 or COV2-2050 LALA-PG at the time of challenge (0 dpi) (mean ± standard error of the mean (SEM); n = 3-4, two experiments). (**D**) Neutralizing titers in serum of indicated groups at 0 dpi as measured by FRNT (mean ± SEM; n = 6, two experiments). (**E**) Weight change following COV2-2050 administration (mean ± SEM; n = 4-6, two experiments: two-way ANOVA with Sidak’s post-test: ns not significant, * *P* < 0.05, **** *P* < 0.0001; comparison to the isotype control mAb-treated group). (**F**) Weight change following COV2-2050 LALA-PG (mean ± SEM; n = 4-6, two experiments: two-way ANOVA with Sidak’s post-test: ns not significant, ** *P* < 0.01, **** *P* < 0.0001; comparison to the isotype control mAb-treated group). (**G**) Correlation analyses comparing COV2-2050 or COV2-2050 LALA-PG serum concentrations (day 0 (D0)) plotted against weight change (D+7) (n = 4-6, two experiments; Pearson’s correlations: COV2-2050, *P* = 0.0026; COV2-2050 LALA-PG, *P* = 0.0006). (**H**) Correlation analyses comparing COV2-2050 neutralizing titers in serum (D0) plotted against weight change (D+7) (n = 6-8, three experiments; Pearson’s correlation: COV2-2050, *P* = 0.0008). (**I**) Viral RNA levels at 7 dpi in the lung as determined by qRT-PCR (n = 4-6, two experiments: one-way ANOVA with Turkey’s post-test: *** *P* < 0.001, **** *P* < 0.0001, comparison to the isotype control mAb-treated group). (**J**) Correlation analyses comparing COV2-2050 or COV2-2050 LALA-PG serum concentrations (D0) plotted against lung viral titer (D+7) (n = 4-6, two experiments; Pearson’s correlation: COV2-2050, *P* = 0.0022; COV2-2050 LALA-PG, *P* = 0.0095). (**K**) Correlation analyses comparing serum neutralizing titers (D0) plotted against lung viral titer (D7) (n = 6-8, two experiments; Pearson’s correlation calculation for COV2-2050, *P* = 0.0006).

Effects on SARS-CoV-2-induced weight loss were comparable between COV2-2050 or COV2-2050 LALA-PG at all doses with a loss of protection observed only at the 1.6 μg (0.08 mg/kg) dose (**Fig 1E-F**). Levels of COV2-2050 correlated with clinical protection (**Fig 1G-H**), with a minimum serum neutralizing titer (NT_50_) of 1:104 and concentration of 212 ng/mL required to prevent weight loss in this stringent challenge model. Although weight loss was prevented at a dose of 1.6 μg per animal, a higher 8 μg (0.4 mg/kg) dose was required to reduce viral burden in the lung at 7 days post-infection (dpi) (**Fig 1I**). Levels of COV2-2050 also correlated inversely with SARS-CoV-2 RNA in the lung (**Fig 1J-K**) with a minimum neutralizing titer of 1:381 and mAb serum concentration of 851 ng/mL required to reduce viral infection at 7 dpi. Notably, differences in viral burden were not observed at several different doses of COV2-2050 and COV2-2050 LALA-PG treatment. Importantly, no evidence of antibody-dependent enhancement of clinical disease or viral burden (Lee et al., 2020; Taylor et al., 2015) was detected even when levels were below the protective dose. Thus, the Fc-dependent effector functions of COV2-2050 are dispensable for clinical and virological protection in K18-hACE2 transgenic mice when a potently neutralizing antibody is administered as prophylaxis.

### Fc effector functions enhance the therapeutic activity of neutralizing antibodies

Although we did not detect a requirement for Fc effector functions when COV2-2050 was administered as prophylaxis, we re-evaluated this observation in the setting of post-exposure administration. We inoculated K18-hACE2 transgenic mice with SARS-CoV-2 by the intranasal route and then delivered a single 200 μg (10 mg/kg) dose of COV2-2050 or COV2-2050 LALA-PG mAbs at 1 (D+1) or 2 (D+2) dpi by intraperitoneal injection. Whereas passive transfer of the intact COV2-2050 prevented weight loss compared to isotype control mAb-treated animals, this protection was lost in animals treated with the COV2-2050 LALA-PG variant (**Fig 2A**). The differences in protection were not due to disparate levels of intact COV2-2050 and COV2-2050 LALA-PG variants, as equivalent amounts were detected in serum at 0 and 4 dpi (**Fig 2B**). Treatment at 1 dpi with COV2-2050, but not COV2-2050 LALA-PG, reduced SARS-CoV-2 viral RNA levels in the lung at 4 and 8 dpi substantially as measured by qRT-PCR (**Fig 2C-D**). In contrast, D+2 treatment with either intact COV2-2050 or COV2-2050 LALA-PG did not reduce viral RNA levels in the lung when compared to isotype-treated controls. However, COV2-2050 and COV2-2050 LALA-PG mAbs both reduced levels of infectious virus in the lung at 4 dpi (**Fig 2E**). Given the disparity in viral RNA levels, this effect might be due in part to *ex vivo* neutralization after lung tissue homogenization, as reported for other respiratory viruses (Subbarao et al., 2004; Wells et al., 1981). Although D+2 treatment did not reduce viral RNA levels in the lung, K18-hACE2 mice receiving intact COV2-2050 at D+2, but not COV2-2050 LALA-PG, showed functional improvement in pulmonary mechanics including inspiratory capacity, respiratory resistance, elastance, tissue damping, and compliance (**Fig 2F**). These results correlated with a smaller downward deflection of the pressure-volume loop and improved lung compliance and distensibility (**Fig 2G**).

**Figure 2.**
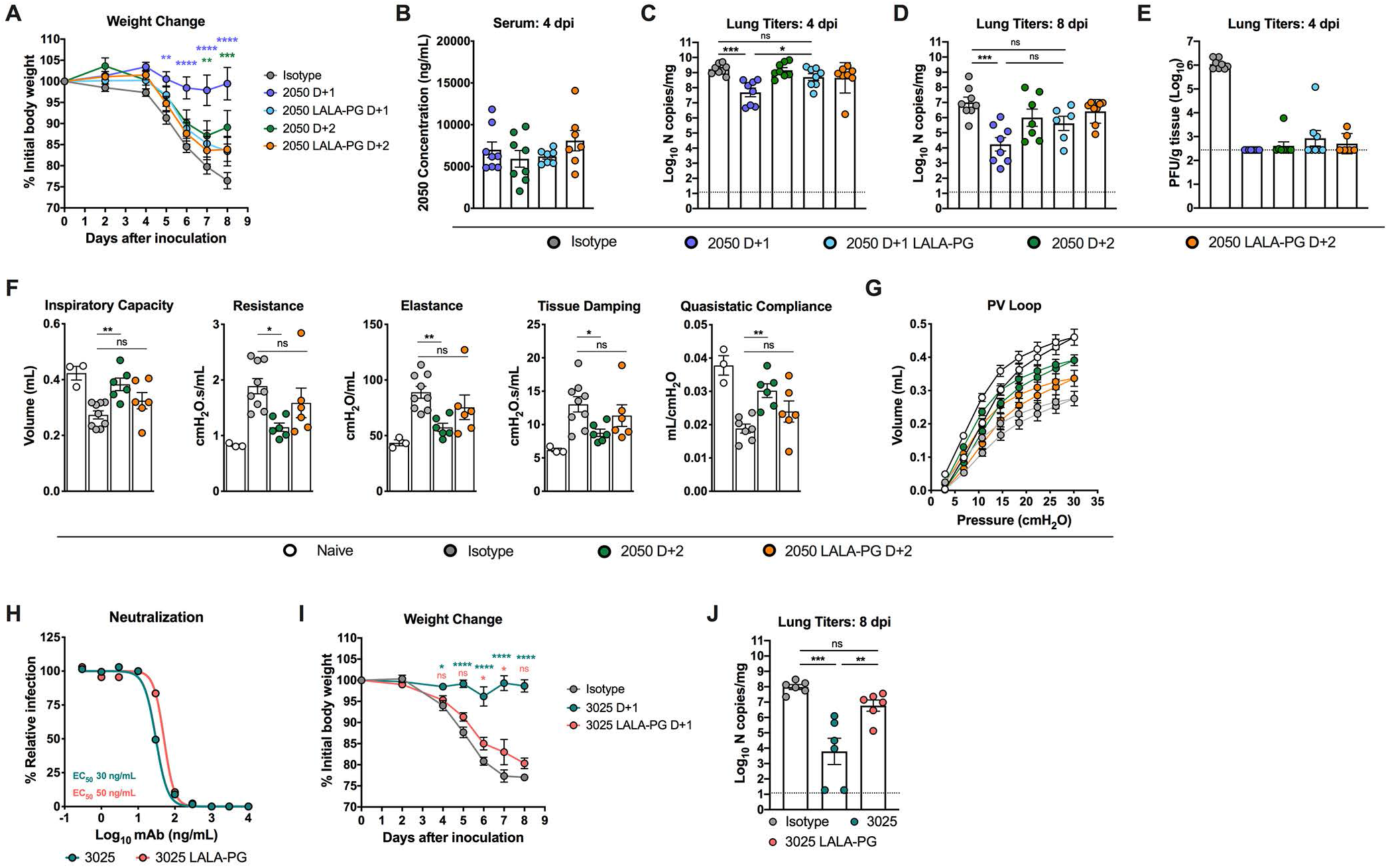
Fc effector functions enhance the therapeutic activity of a neutralizing antibody against SARS-CoV-2 in K18-hACE2 transgenic mice. A-F. Eight-week-old female and male K18-hACE2 transgenic mice were inoculated by the intranasal route with 10^3^ PFU of SARS-CoV-2. At 1 (D+1) or 2 (D+2) dpi, mice were given a single 200 μg dose of COV2-2050 or COV2-2050 LALA-PG by intraperitoneal injection. Naïve animals were mock-infected with sterile PBS. (**A**) Weight change (mean ± SEM; n = 8-10, three experiments: two-way ANOVA with Sidak’s post-test: * *P* < 0.001, *** *P* < 0.001, **** *P* < 0.0001; comparison to the isotype control mAb-treated group). (**B**) Serum concentrations (ng/mL) of COV2-2050 or COV2-2050 LALA-PG at 4 dpi (mean ± SEM; n = 8, two experiments). (**C-D**) Viral RNA levels at 4 and 8 dpi in the lung as determined by qRT-PCR (n = 8-10, three experiments: one-way ANOVA with Turkey’s post-test: ns not significant, * *P* < 0.05, *** *P* < 0.001, **** *P* < 0.0001, comparison to the isotype control mAb-treated group). (**E**) Infectious virus as measured by plaque assay at 4 dpi in the lung (n = 8, two experiments). (**F**) Parameters of respiratory mechanics: inspiratory capacity, resistance, elastance, tissue damping, and quasistatic compliance measured at 8 dpi (n = 3-6, two experiments: two-way ANOVA with Turkey’s post-test: ns not significant; * *P* < 0.05, ** *P* < 0.01, comparison to the isotype control mAb-treated group). (**G**) Pressure volume loops (n = 3-6, two experiments). (**H**) Anti-SARS-CoV-2 mAbs (COV2-3025 and COV2-3025 LALA-PG) were incubated with 10^2^ FFU of SARS-CoV-2 for 1 h at 37°C followed by addition of mAb-virus mixture to Vero E6 cells. Wells containing mAb were compared to those without mAb to determine relative infection. One experiment of two, with similar results, is shown. The mean of two technical replicates is shown. **I-J.** Eight-week-old male K18-hACE2 transgenic mice were inoculated by the intranasal route with 10^3^ PFU of SARS-CoV-2. At 1 (D+1) dpi, mice were given a single 200 μg dose of COV2-3025 or COV2-3025 LALA-PG by intraperitoneal injection. (**I**) Weight change (mean ± SEM; n = 6, two experiments: two-way ANOVA with Sidak’s post-test: * *P* < 0.05, **** *P* < 0.0001; comparison to the isotype control mAb-treated group). (**J**) Viral RNA levels at 8 dpi in the lung as determined by qRT-PCR (n = 6, two experiments: one-way ANOVA with Turkey’s post-test: ns not significant, ** *P* < 0.01, *** *P* < 0.001, comparison to the isotype control mAb-treated group).

To corroborate these results, we tested the therapeutic activity of additional neutralizing human mAbs that bind the RBD on the S protein, COV2-3025, COV2-2072, and COV2-2381 (Zost et al., 2020b), and compared them to corresponding LALA-PG or LALA variants of each mAb. For COV2-2381, we used a LALA variant of COV2-2381, since this version also is commonly used to minimize Fc effector functions (Hessell et al., 2007; Sapparapu et al., 2016). The LALA form lacks the P329G mutation, which is needed to fully abolish binding to all mouse FcγRs (*e.g*., mouse FcγRII) and prevent complement deposition (Arduin et al., 2015; Lo et al., 2017; Sondermann et al., 2000). COV2-3025, COV2-2072, and COV2-2381 neutralized SARS-CoV-2 equivalently compared to their respective LALA-PG or LALA variants (**Fig 2H, S1A, and S1E**) and introduction of the LALA mutation abolished binding to mouse FcγRI and FcγRIV (**Fig S1B**). Administration of intact COV2-3025 at D+1 recapitulated the same pattern of protection observed with COV2-2050 treatment: COV2-3025 LALA-PG failed to protect mice from weight loss or reduce viral titers compared to the intact COV2-3025 (**Fig 2I-J**).

In comparison, COV2-2381 LALA partially protected against weight loss, and viral burden was reduced in mice receiving either COV2-2381 or COV2-2381 LALA compared to isotype-treated controls, although we noted a trend towards lower levels of viral RNA in animals receiving the intact COV2-2381 (**Fig S1C-D**). Administration of COV2-2072 LALA-PG at D+1 partially protected against SARS-CoV-2-induced weight loss compared to the isotype-treated controls, but failed to reduce viral titers, unlike the intact COV2-2072 (**Fig S1F-G**). The relative differences in therapeutic capacity of COV2-2050 LALA-PG and COV2-3025 LALA-PG compared to COV2-2381 LALA and COV2-2072 LALA-PG antibodies could be due to differential neutralizing activity *in vivo*, disparate affinity for FcγR or complement, or because these mAbs, which recognize distinct epitopes on RBD (Zost et al., 2020b), variably engage immune effector molecules based on their orientation of binding to the virion (Renner et al., 2018). Regardless, these results indicate that intact Fc effector functions of neutralizing antibodies are needed for optimal therapeutic activity to mitigate clinical disease caused by SARS-CoV-2 infection. Notwithstanding this point, some antibodies (*e.g*., COV2-2381 and COV2-2072) might confer therapeutic protection in the absence of Fc effector functions at least at early time points post-infection, possibly due to differences in neutralizing activity *in vivo.*

### Fc effector functions of a neutralizing antibody modulate the immune responses to SARS-CoV-2 infection

An excessive pro-inflammatory host response to SARS-CoV-2 infection is hypothesized to contribute to pulmonary pathology and severe COVID-19 (Giamarellos-Bourboulis et al., 2020). Given that COV2-2050-mediated improvement in pulmonary function was not associated with reduced viral RNA levels in the lung when given at D+2, we speculated that the intact mAb might modulate immune cell and inflammatory responses. Consistent with our previous study (Winkler et al., 2020), analysis of hematoxylin and eosin-stained lung sections from isotype mAb-treated mice at 8 dpi showed perivascular and parenchymal immune cell accumulation with accompanying edema and lung consolidation (**Fig 3A**). Animals receiving COV2-2050 at D+1 showed markedly reduced lung inflammation but this protection was lost in mice receiving COV2-2050 LALA-PG at D+1 or D+2. While COV2-2050 treatment at D+2 did not completely reduce pathology, immune cell infiltration was more focal in nature with patches of normal-appearing airspaces throughout the lung.

**Figure 3.**
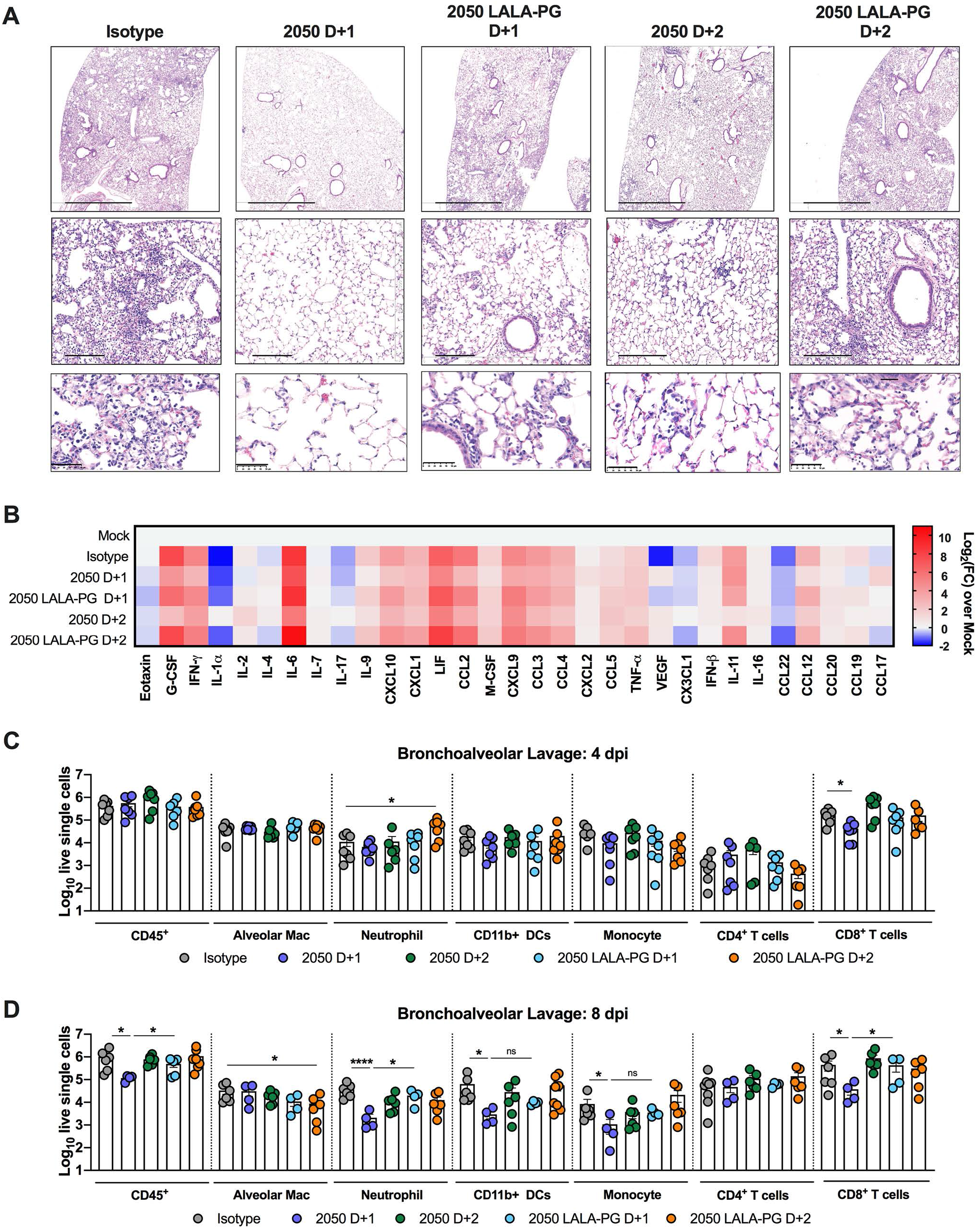
Fc effector functions of a neutralizing antibody modulate the immune responses to SARS-CoV-2 infection. Eight-week-old female and male K18-hACE2 transgenic mice were inoculated by the intranasal route with 10^3^ PFU of SARS-CoV-2. At 1 (D+1) or 2 (D+2) dpi, mice were given a single 200 μg dose of COV2-2050 or COV2-2050 LALA-PG by intraperitoneal injection. (**A**) Hematoxylin and eosin staining of lung sections at 8 dpi. Images show low- (top; scale bars, 1 mm), medium-power (middle; scale bars, 200 μm), and high-power (bottom; scale bars, 50 μm). Representative images from n = 3 per group. (**B**) Heat-maps of cytokine levels as measured by multiplex platform in lung tissue of SARS-CoV-2-infected mice at 8 dpi. For each cytokine, fold-change was calculated compared to mock-infected animals and log2(fold-change) was plotted in the corresponding heat-map (three experiments, n = 5 per group except naïve (n = 2), associated statistics are reported in **Figure S2**). **C-D.** Flow cytometric analysis of immune cells from BAL fluid harvested at (**C**) 4 and (**D**) 8 dpi post-SARS-CoV-2 infection (two experiments, n = 4-8 per group; bars represent the mean. One-way ANOVA with Dunnett’s test; * *P* < 0.05; **** *P* < 0.0001.) Gating scheme shown in **Fig S3**.

Measurements of pro-inflammatory cytokine and chemokines in the lung at 8 dpi showed decreased levels of CXCL10, G-CSF, IL-6, IFN-γ, CCL2, CCL3, CCL4, CCL19, and CXCL1 following both D+1 and D+2 COV2-2050 treatment, which did not occur in animals receiving COV2-2050 LALA-PG (**Fig 3B and S2**). Cellular analysis of bronchoalveolar lavage (BAL) fluid at 4 dpi (**Fig S3**, using the gating scheme of (Misharin et al., 2013)) showed significantly reduced numbers of CD8^+^ T cells with a trend towards fewer monocytes after administration of intact COV2-2050 at D+1. Differences in BAL fluid cell numbers were not observed at 4 dpi when treatment with COV2-2050 or COV2-2050 LALA-PG was started at D+2 (**Fig 3C**). However, at 8 dpi, we observed a significant reduction in the numbers of CD45^+^ cells, neutrophils, and CD8^+^ T cells in mice treated at D+1 with COV2-2050 compared to COV2-2050 LALA-PG (**Fig 3D**), consistent with histopathological findings (**Fig 3A**). However, the only difference in immune cell infiltrates in BAL fluid at D+2 was a reduced number of alveolar macrophages in animals receiving COV2-2050 compared to COV2-2050 LALA-PG (**Fig 3D**). Given that viral RNA levels were equivalent in mice receiving COV2-2050 and COV2-2050 LALA-PG at D+2 (**Fig 2C and E**) and only small differences in BAL cell number were observed, Fc effector engagement may shape inflammatory immune responses independently of effects on viral burden or the number and type of immune cells recruited.

### Distinct transcriptional signatures in the lung after treatment with COV2-2050 or COV2-2050 LALA-PG

To interrogate further the impact of Fc effector functions on protection, we performed RNA sequencing of lung homogenates at 8 dpi in mice receiving an isotype control mAb, COV2-2050, or COV2-2050 LALA-PG mAbs at 1 (D+1) or 2 (D+2) and compared the results to those from samples from naïve mice. Principal component analysis (PCA) revealed distinct transcriptional signatures associated with SARS-CoV-2 infection when compared to mock-infected naïve animals. Treatment with intact COV2-2050 demonstrated a clear transcriptional shift towards the naive group, whereas the profiles from COV2-2050 LALA-PG-treated mice were more similar to isotype control mAb-treated animals (**Fig 4A**). Indeed, only 91 differentially expressed genes (DEGs) were identified when comparing the isotype control mAb to the COV2-2050 LALA-PG D+1 group, whereas 2056 and 1975 DEGs were detected when comparing COV2-2050 D+1 to isotype control and COV-2050 LALA-PG D+1, respectively (**Fig 4B**). A similar large number of DEGs was observed in the lungs at 8 dpi from mice treated at D+2 with COV-2050 or COV2-2050 LALA-PG (**Fig 4B**). Gene ontology analysis of the top downregulated genes comparing COV2-2050 to COV2-2050 LALA-PG showed immune gene clusters including cytokine-mediated signaling (*e.g*., *Il10ra, Il15, Il17ra, Socs1, and Jak3*), type I IFN signaling (*e.g*., *Ifnar2, Stat2, Ifit1, Ifit2, Ifit3, and Irf7*), and leukocyte chemotaxis (*e.g*., *Ccl2, Cxcl10, and Ccl7*) as well as genes involved in cell proliferation (*e.g*., *Egfr, Fgfr1, Fosl1, Myc, and Cdkn2b*) and metalloproteinase-mediated extracellular matrix organization (*e.g*., *Adam15, Adam19, Col1a, Mmp14, and Itgam*) (**Fig S4 and Table S1**).

**Figure 4.**
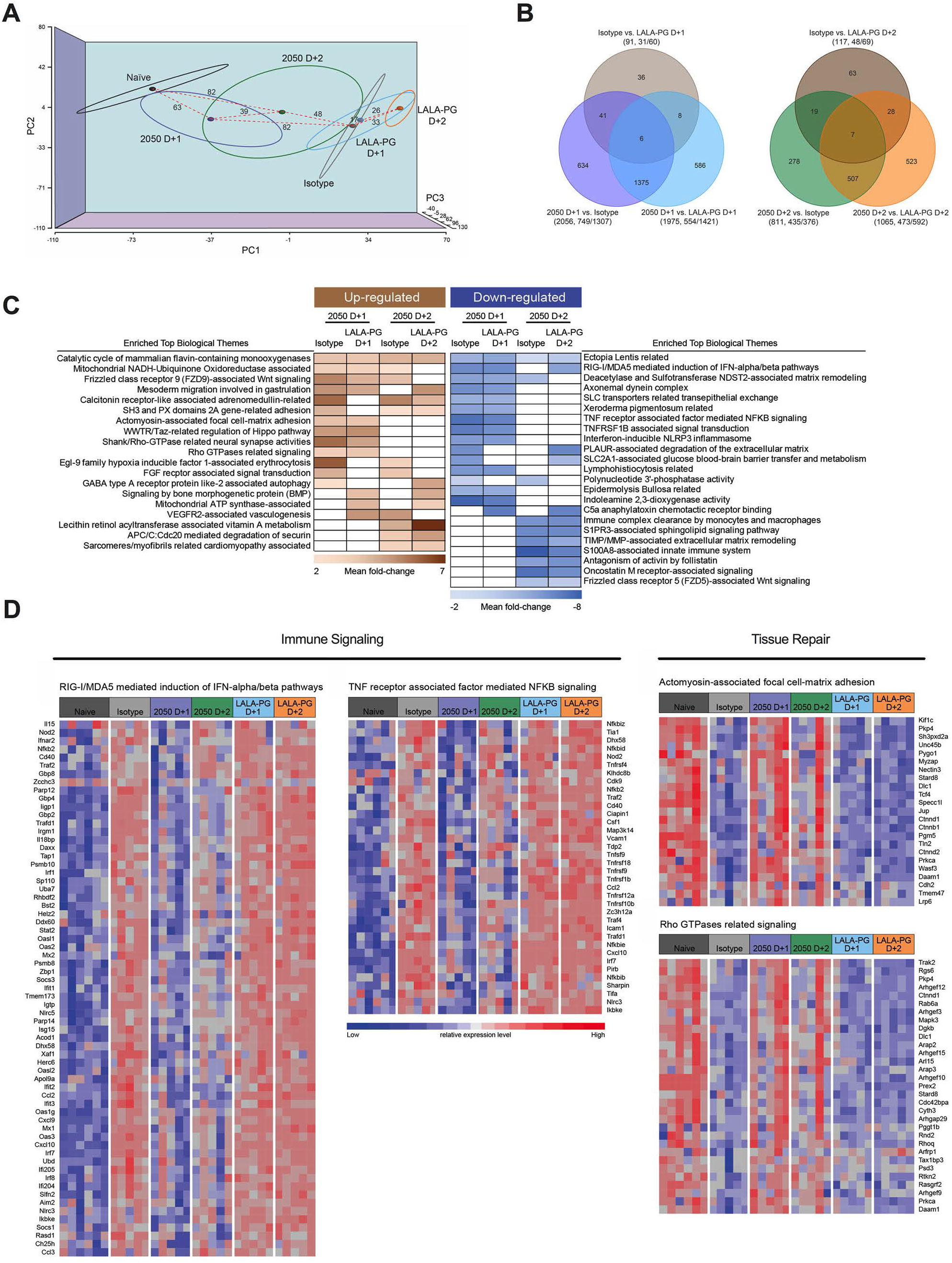
Distinct transcriptional signatures in the lung are associated with COV2-2050 with intact Fc effector functions. RNA sequencing analysis from the lung homogenates of naive K18-hACE2 mice or mice infected with SARS-CoV-2 infection at 8 dpi. At 1 (D+1) or 2 (D+2) dpi, mice were given a single 200 μg dose of COV2-2050 or COV2-2050 LALA-PG by intraperitoneal injection. (**A**) Three-dimensional map from principal component analysis (PCA) of the RNA-seq data in the study. The PCA has been performed using 12,157 unique genes with count per million reads ≥ 1 in at least 5 of the study samples (n = 31). Each group is represented by an ellipse and the color-matched solid circle, which is the centroid of each group. The size of the ellipse is the centroid with one standard deviation. The dashed red lines with numbers indicate the spatial distance between centroids of the 6 groups, which is calculated by using the three-dimensional coordinates for the centroids. (**B**) Venn diagrams of overlapping genes identified in differential expression analysis when comparing isotype control, COV2-2050 D+1, and COV2-2050 LALA-PG D+1 or isotype control, COV2-2050 D+2, and COV2-2050 LALA-PG D+2. Numbers in the parenthesis under each comparison indicate the number of differentially-expressed genes (fold-change >2 at *P* < 0.05) followed by the proportion that are up or down-regulated. (**C**) The significantly enriched biological themes defined by a novel pathway analysis tool ‘CompBio’ comparing treatments with isotype control mAb, COV2-2050 (D+1 and D+2), and COV2-2050 LALA-PG (D+1 and D+2). Only those themes enriched in at least two comparisons are displayed. These themes either are upregulated (brown color) or downregulated (blue color) in the COV2-2050-treated group (at D+1 or D+2) when compared to the isotype control or COV2-2050 LALA-PG-treated groups. The scaled color blocks represent the mean fold-change of enriched genes with an enrichment score of 10 or greater in the comparison. (**D**) Heatmaps of selected relevant biological themes (RIG-I/MDA-5 mediated signaling, TNF receptor-associated signaling, actinomyosin cell adhesion, Rho GTPases related signaling) enriched in COV2-2050 D+1 versus isotype control or COV2-2050 LALA-PG D+1. Genes shown are common in the pair of comparisons having an enrichment score of 100 or greater.

We next performed CompBio analysis (v2.0, PercayAI), as we did previously (Adamo et al., 2020; Gehrig et al., 2019), to identify biological themes uniquely enriched in the COV2-2050-treated groups compared to the isotype control and COV2-2050 LALA-PG-treated groups (**Fig 4C, Table S2**). Pathways unique to the COV2-2050 D+1 treatment group compared to the isotype-treated group included genes involved in actinomyosin-associated cell adhesion (*e.g*., *Kif1c, Ctnnd1, Nectin3, Unc45b, and Prkca*) and Rho GTPase signaling (*e.g*., *Rhoq, Mapk3, Prkca, and Cdc42bpa*), processes that are typically associated with wound repair programs (Verboon and Parkhurst, 2015) (**Fig 4D**). The expression pattern of these gene sets in the COV2-2050 D+1 treatment was similar to naïve animals, suggesting that the intact antibody limited virus-induced perturbations in transcription. Pathways uniquely downregulated in the COV2-2050 D+1 group included genes involved in type I IFN and NFκB-dependent signaling (*e.g., Irf7, Stat2, Nfkb2, Bst2, Isg15, and Ikbk3*), which may in part be due to the lower levels of viral RNA detected (**Fig 2C-D and 4D**).

Pathways that were downregulated in the D+2 COV2-2050-treated group compared to the isotype control or D+2 COV2-2050 LALA-PG-treated animals included S100A8-associated innate immune signaling (*e.g*., *Reg3g, Saa3, Itgma, Mmp8, and S100a8*), oncostatin M receptor associated signaling (*e.g*., *Il6, Osmr, Csf1, and Socs3*), and extracellular matrix remodeling (*e.g*., *Adamts15, Col5a1, Vcam1, and Lama4*) (**Fig S5, Table S2**). Thus, COV2-2050 treatment at D+2 and the resultant Fc-dependent effector responses may differentially modulate neutrophil activation, gp130 signaling, and tissue damage due to matrix metalloproteinase activation. Of note, we also identified diminished expression of genes involved in immune complex clearance following COV2-2050 versus COV2-2050 LALA-PG administration at D+2, which may reflect a negative feedback loop of this pathway (Daëron and Lesourne, 2006). Collectively, our results with intact COV-2050 suggest that Fc engagement by FcγR and/or complement can induce distinct and protective transcriptional programs in the SARS-CoV-2-infected lung depending on the timing of therapy.

### Monocytes are required for optimal therapeutic activity of neutralizing mAb

Fc engagement of FcγRs on innate immune cells including macrophages, monocytes, NK cells, and neutrophils can lead to antibody-dependent cellular cytotoxicity (ADCC) or phagocytosis (ADCP), modulation of inflammatory mediators produced by these cells, and enhanced adaptive immunity (Bournazos et al., 2020; Lu et al., 2018). To determine which innate immune cells contribute to the antibody-mediated protection observed *in vivo*, we depleted Ly6^hi^ classical monocytes (anti-CCR2) (**Fig 5A and S6**), natural killer (NK) cells (anti-NK1.1) (**Fig 5B and S6**), or neutrophils (anti-Ly6G) (**Fig 5C and S6**) in combination with COV2-2050 treatment at D+1. Depletion of neutrophils or NK cells did not affect SARS-CoV-2 pathogenesis in the presence of COV2-2050 or the isotype control mAb (**Fig 5B and C**). However, when monocytes were depleted, COV2-2050 failed to prevent the weight loss phenotype seen in non-depleted, COV2-2050-treated mice (**Fig 5A**). Moreover, monocyte depletion in the setting of COV2-2050 therapy was associated with greater immune cell infiltration and lung damage (**Fig 5D**). Indeed, expression of *Ccl2, Cxcl10*, and *Il6* were increased following COV2-2050 treatment and monocyte depletion compared to treatment with COV2-2050 or an isotype control mAb (**Fig 5E**). The monocyte depletion phenotype, however, was not associated with changes in viral RNA levels at 8 dpi (**Fig 5F**). This result suggests that monocytes are a key immune cell type that mediates therapeutic antibody-dependent protection through a mechanism that at least is partially independent of viral clearance.

**Figure 5.**
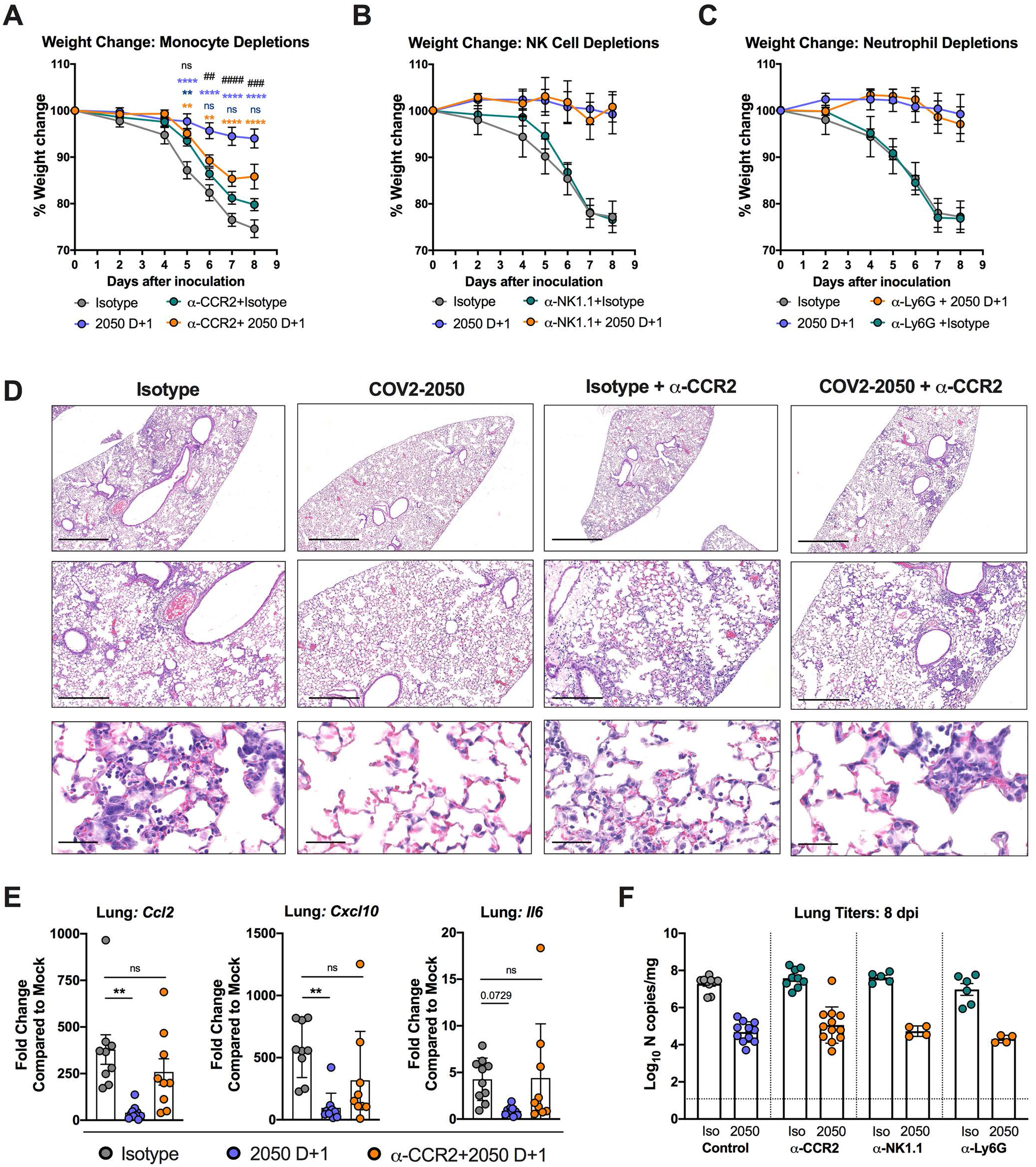
Monocytes are necessary for protection following mAb therapy. **A-D**. Eight-week-old female or male K18-hACE2 transgenic mice received the cell depleting antibodies anti(α)-CCR2 (50 μg/dose) (**A**), α-NK1.1 (200 μg/dose) (**B**), or α-Ly6G (250 μg/dose) (**C**) or corresponding isotype controls at D-1, D+1, D+3, D+5, and D+7 relative to SARS-CoV-2 infection. At D0, animals were inoculated by the intranasal route with 10^3^ PFU of SARS-CoV-2. At 1 (D+1) or 2 (D+2) dpi, mice were administered a single 200 μg dose of COV2-2050 by intraperitoneal injection. (**A-C**) Weight change (mean ± SEM; n = 8-12, 2-3 experiments: two-way ANOVA with Sidak’s post-test: * *P* < 0.001, *** *P* < 0.001, **** *P* < 0.0001; comparison to the isotype control mAb-treated group). (**D**) Hematoxylin and eosin staining of lung sections at 8 dpi. Images show low- (top; scale bars, 1 mm), medium-power (middle; scale bars, 250 μm), or high-power (bottom; scale bars, 50 μm). Representative images from n = 3 per group. (**E**) Fold change in gene expression of indicated cytokines and chemokines in lung homogenates as determined by RT-qPCR, normalized to *Gapdh*, and compared to naïve controls (three experiments, n = 8-12 per group, One-way ANOVA with Dunnett’s test; ** *P* < 0.01). Dotted lines indicate the mean level of cytokine or chemokine transcript in naïve mice. (**F**) Viral RNA levels at 8 dpi in the lung as determined by qRT-PCR (n = 8-12, 2-3 experiments).

### Fc effector functions enhance the therapeutic activity of neutralizing antibodies against SARS-CoV-2 in Syrian hamsters

To validate the requirement for Fc effector functions in optimal therapeutic antibody protection, we used a golden Syrian hamster model. This animal model supports viral replication with corresponding weight loss and interstitial pneumonia (Rosenke et al., 2020; Sia et al., 2020) and has been used to evaluate mAb-based therapies and vaccines (Liu et al., 2020; Rogers et al., 2020; Tostanoski et al., 2020) against SARS-CoV-2. We inoculated seven-month-old hamsters with SARS-CoV-2 (5 × 10^5^ PFU, 2019 n-CoV/USA_WA1/2020 strain) by the intranasal route and then delivered a single 1 mg (10 mg/kg) dose of COV2-2050 or COV2-2050 LALA-PG mAbs at 1 dpi (D+1) by intraperitoneal injection. Consistent with observations in mice, passive transfer of intact COV2-2050 prevented weight loss compared to isotype mAb-treated animals at 5 and 6 dpi, and this protection was lost in animals treated with the COV2-2050 LALA-PG variant (**Fig 6A**). Furthermore, hamsters treated with intact COV2-2050, but not COV2-2050 LALA-PG, showed a reduction in viral RNA levels at 6 dpi (**Fig 6B**). The improved viral burden with COV2-2050 was associated with lower levels of the inflammatory mediators *Cxcl10, Ccl2, Ccl3, Ccl5*, and *Ifit3* (**Fig 6C**). Thus, therapeutic efficacy following neutralizing mAb administration also depends on Fc interactions in a second, relevant small animal model.

**Figure 6.**
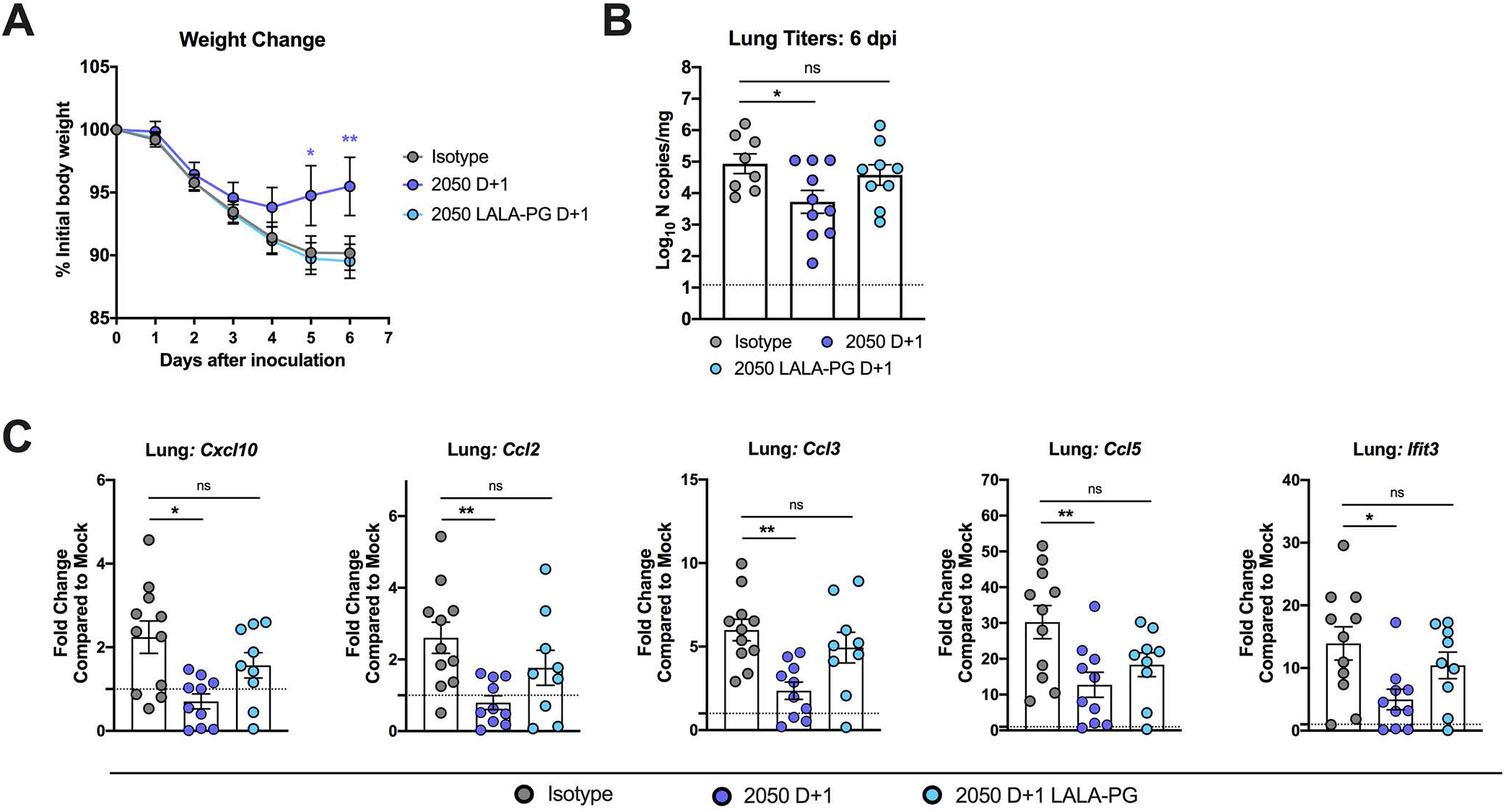
Fc effector functions enhance the therapeutic activity of neutralizing antibodies against SARS-CoV-2 in Syrian hamsters. **A-C** Seven-month-old female Syrian hamsters were inoculated by the intranasal route with 5 × 10^5^ PFU of SARS-CoV-2. At 1 dpi (D+1), hamsters were given a single 1 mg dose of COV2-2050 or COV2-2050 LALA-PG by intraperitoneal injection. (**A**) Weight change (mean ± SEM; n = 8-10, two experiments: two-way ANOVA with Sidak’s post-test: * *P* < 0.05, ** *P* < 0.01; comparison to the isotype control mAb-treated group). (**B**) Viral RNA levels at 6 dpi in the lung as determined by qRT-PCR (n = 8-10, two experiments: one-way ANOVA with Turkey’s post-test: ns not significant, * *P* < 0.05, comparison to the isotype control mAb-treated group). (**C**) Fold change in gene expression of indicated cytokines and chemokines in lung homogenates as determined by RT-qPCR, normalized to *Rpl18*, and compared to naïve controls (two experiments, n = 8-10 per group, One-way ANOVA with Dunnett’s test; * P < 0.05; ** P < 0.01). Dotted lines indicate the mean cytokine or chemokine transcript levels in naïve hamsters.

## DISCUSSION

Neutralizing monoclonal antibodies against SARS-CoV-2 are a promising option for the treatment of COIVD-19 and have demonstrated therapeutic efficacy in a variety of pre-clinical models (Abraham, 2020; Marovich et al., 2020). Antibody programs have reported encouraging results in clinical trials (Chen et al., 2020), and recently, the FDA granted Emergency Use Authorization to the anti-SARS-CoV-2 S mAb bamlanivimab for newly diagnosed, mild-to-moderately ill, high-risk patients. However, the mechanisms of protection *in vivo* have not been fully investigated and are assumed to be directly related to the neutralizing capacity of the mAb. In this study, we examined the contributions of Fc effector functions for clinical and virological protection in murine and hamster models of SARS-CoV-2 infection and pathogenesis. Whereas intact Fc effector functions were not required when neutralizing mAbs were given as prophylaxis, for several anti-RBD human mAbs tested in mice, a functional Fc region was required for optimal protection as post-exposure therapy. This enhanced therapeutic efficacy of intact COV2-2050 and COV2-3025 compared to their LALA-PG IgG loss-of-function Fc region variants correlated with decreased viral burden, improved pulmonary function, diminished inflammatory responses, and preserved tissue repair processes. In mice, cell depletion studies identified monocytes as a key immune cell type for antibody-dependent clinical protection.

Fc effector function interactions were not required when neutralizing mAbs were administered as prophylaxis, suggesting their mechanism of protection in a pre-exposure setting depends largely on the neutralizing capacity of the antibody to prevent initial viral infection and limit dissemination. In the stringent K18-hACE2 transgenic model of SARS-CoV-2 pathogenesis (Golden et al., 2020; Winkler et al., 2020), we defined minimum serum neutralizing titers (NT_50_) and concentrations of 1:104 and 212 ng/ml for the prevention of weight loss and 1:381 and 851 ng/ml for reduction of viral burden in the lung. Establishing serum correlates of protection for mAb- and vaccine-based therapies in pre-clinical models is an important first step for translation and analysis of human studies. Levels of neutralizing antibody required for protection against disease in humans might be lower due to the slower kinetics of disease pathogenesis and/or a longer half-life of the antibody. Lastly, even at sub-protective doses of intact COV2-2050 (0.08 mg/kg), we did not observe evidence of antibody-dependent enhancement (ADE) of infection or immune enhancement, consistent with other SARS-CoV-2 vaccine and antibody treatment studies (Baum et al., 2020; Cao et al., 2020; Halstead and Katzelnick, 2020; Laczkó et al., 2020; Luo et al., 2018; Qin et al., 2006; Rogers et al., 2020).

In contrast to prophylactic administration, Fc effector functions were required for optimal efficacy when neutralizing mAbs were administered as post-exposure therapy. The treatment window to reduce viral RNA levels in the lung in the stringent K18-hACE2 transgenic model was essentially one day, suggesting that antibody therapy optimally should be given prior to the peak of viral replication, which is 2 dpi in this model (Winkler et al., 2020). The narrow therapeutic window for reducing viral load we observed could be related to the stoichiometry of antibody binding required for virus neutralization (Pierson and Diamond, 2015), such that when too much viral antigen is present in the lung, neutralization is limited. Against this hypothesis, infectious virus levels were neutralized and equivalently low at 4 dpi in mice treated with intact or LALA-PG versions of antibodies at 1 or 2 dpi, although it is possible that this effect is due to *ex vivo* neutralization of virus after tissue homogenization (Subbarao et al., 2004; Wells et al., 1981). More likely, the Fc effector functions of neutralizing antibodies serve to clear SARS-CoV-2 infected cells either directly by engagement of FcγR on myeloid cells and subsequent ADCP or ADCC or through enhanced antigen presentation and induction of antigen-specific CD8^+^ T cell responses (Bournazos et al., 2020). Although the precise mechanism awaits delineation, our data suggests that in the post-exposure therapeutic setting, the neutralizing activity of most RBD-specific antibodies may no longer be sufficient for optimal protection, and additional mechanisms mediated by Fc effector functions are required. We acknowledge that some anti-RBD mAbs with superior neutralizing activity *in vivo* may confer protection in the absence of Fc effector functions at least at early time points post-infection. However, at later time points, once infection is established and high amounts of viral RNA are produced in the lung, Fc effector functions of even ‘elite’ neutralizing mAbs may be needed for immune modulation and the most favorable clinical outcome.

With the lower levels of viral RNA in lungs of mice treated at 1 dpi with COV2-2050 but not COV2-2050 LALA-PG, we also found reduced levels of pro-inflammatory cytokines, immune cell infiltration, and expression of genes downstream of type I IFN and NFκB-dependent signaling pathways. Transcriptional signatures associated with COV2-2050 therapy also included enrichment of gene sets involved in tissue repair processes. This same pattern of expression was observed in naïve mice, suggesting that Fc effector functions of antibodies may limit virus-induced perturbations and maintain certain reparative homeostatic transcriptional programs. Remarkably, administration of neutralizing mAb at 2 dpi still improved clinical outcome and pulmonary function in an Fc-dependent manner without substantive reduction in viral RNA levels. The gene signatures uniquely associated with intact COV2-2050 mAb administration at 2 dpi differed from those at 1 dpi, as they showed reduced expression of genes involved in neutrophil activation, IL-6 signaling, and metalloproteinase-mediated extracellular matrix remodeling pathways. Altogether, these results suggest that at least some neutralizing mAbs protect *in vivo* through multiple Fc-dependent mechanisms including viral clearance and modulation of the immune response to enhance resolution of inflammation.

The importance of Fc effector functions for antibody efficacy *in vivo* has been illustrated in other viral models, including HIV, Ebola, West Nile, hepatitis B, chikungunya, and influenza viruses (Bournazos et al., 2020; DiLillo et al., 2016; DiLillo et al., 2014; Fox et al., 2019; Hessell et al., 2007; Li et al., 2017; Liu et al., 2017; Vogt et al., 2011), although the specific cells mediating this protection have been characterized in relatively few cases (Fox et al., 2019; He et al., 2017). Our immune cell depletion studies showed that monocytes, but not neutrophils or NK cells were necessary for mAb-dependent clinical protection and diminished pro-inflammatory cytokine responses following SARS-CoV-2 infection in mice, although monocyte depletion did not affect viral burden. Circulating CCR2^+^ monocytes can differentiate into a different myeloid cell subsets including interstitial macrophages and monocyte-derived dendritic cells following migration into the lung (Jakubzick et al., 2017). The phenotype of these myeloid cells in the lung can vary widely ranging from TNF/iNOS producing DCs (Tip-DCs) (Serbina et al., 2003) to interstitial macrophages involved in tissue repair, and myeloid cell dysregulation has been linked to severe COVID-19 in humans (Schulte-Schrepping et al., 2020). Thus, Fc effector engagement on monocyte-derived cells in the lung may not directly impact viral clearance, but rather limit immunopathology through reprograming of the inflammatory response. Consistent with this idea, administration of intact COV2-2050 but not COV2-2050 LALA-PG improved clinical outcome and mitigated inflammation without decreasing viral RNA levels in the lung. Notwithstanding these points, it remains to be determined which cell type is responsible for the Fc-effector function dependent reductions in viral burden we observed with treatment at D+1; this could occur through phagocytosis of infected cells by alveolar macrophages or other myeloid cells that do not express CCR2 (He et al., 2017) or enhanced antigen presentation, accelerated priming of CD8^+^ T cells, and cytolysis of virally infected cells (Bournazos et al., 2020). Engineering of the Fc domain sequences to optimize or possibly enhance effector functions (Bournazos et al., 2020; Lazar et al., 2006; Saunders, 2019) could have implications for the therapeutic activity of neutralizing mAbs against SARS-CoV-2.

Our results highlight the importance of Fc effector functions of antibody in two animal models of SARS-CoV-2 infection. While LALA-PG mutations abrogate binding to both FcγRs and C1q, it remains unknown which of these effector molecules or particular receptors (*e.g*., FcγRI, FcγRIII, or FcγRIV) dominantly associate with loss of anti-SARS-CoV-2 mAb therapeutic activity. Confirmatory studies also are needed in non-human primates, a model that more closely recapitulates human mAb dosing kinetics *in vivo*. Moreover, we used human IgG1 anti-SARS-CoV-2 antibodies in mice and hamsters; FcγR expression patterns on immune cells and human IgG-mouse/hamster FcγR interactions may not fully recapitulate patterns observed with human IgG, human FcγRs, and human cells (Bournazos et al., 2014a). While transgenic mice expressing human FcγRs exist (Smith et al., 2012), they are not on a congenic C57BL/6 background and have not been crossed to the K18-hACE2 transgenic mice. Future studies with human FcγR mice, mouse-adapted strains of SARS-CoV-2 (Leist et al., 2020), and additional human anti-SARS-CoV-2 mAbs may be useful for confirmation of the phenotypes described.

The requirement of Fc effector functions for protection by other anti-SARS-CoV-2 mAbs warrants more study. Indeed, *in vitro* neutralization potency does not uniformly correlate with *in vivo* protection (Schäfer et al., 2020). Our studies used neutralizing antibodies that bind the RBD on S and block ACE2 receptor engagement (Zost et al., 2020b). Other neutralizing mAbs against the N-terminal domain or some classes of non-neutralizing mAbs also may require Fc effector functions for therapeutic activity. Even among anti-RBD mAbs, we observed variation, as clinical protection with COV2-2381 and COV2-2072 was only partially Fc-dependent when administered as post-exposure therapy at D+1, yet virological protection was lost with COV2-2072 LALA-PG, but not COV2-2381 LALA administration. This result could be due to differences in epitope binding or neutralization potency between different anti-RBD antibodies or the angle of engagement and accessibility of the Fc region for C1q or FcγR binding. In conclusion, our study highlights the contributions of Fc effector functions for therapeutic activity of neutralizing mAbs through reductions in viral burden and/or immunopathology. Accordingly, the design of antibody-based combinations against SARS-CoV-2 likely should optimize both neutralization and Fc effector function activities to enhance the window of treatment and provide the greatest virological and clinical protection.

## Supporting information

Table S1 - Gene lists with GO analysis

Table S2 - Gene lists with CompBio analysis

## ACKNOWLEDGEMENTS

This study was supported by contracts and grants from NIH (75N93019C00062, 75N93019C00074, R01 AI157155, U01 AI15181), and the Defense Advanced Research Project Agency (HR001117S0019), Fast Grants (Mercatus Center, George Mason University), the Dolly Parton COVID-19 Research Fund at Vanderbilt University, and the Future Insight Prize (Merck KGaA; to J.E.C). E.S.W. is supported by T32 AI007163, and J.B.C. is supported by a Helen Hay Whitney Foundation postdoctoral fellowship. This work also was funded, in part, by the University of Georgia (UGA) (UGA-001) and Washington University. T.M.R. is supported by the Georgia Research Alliance as an Eminent Scholar. We thank the Pulmonary Morphology Core at Washington University School of Medicine for tissue sectioning and slide preparation. We also thank Joseph Reidy and Andrew Trivette at Vanderbilt Vaccine Center for human antibody sequence analysis and verification, Ron Cobb of Ology Bioservices and Chris Earnhart of the U.S. Joint Program Executive Office for Chemical, Biological, Radiological and Nuclear Defense for providing COV2-2381 and COV2-2381-LALA antibodies, and Rachel Nargi for assistance with mAb purification.

## AUTHOR CONTRIBUTIONS

P.G., S.J.Z., R.E.S., and R.H.C. designed and generated human antibodies. E.S.W., A.L.B., and R.E.C. performed mouse experiments and clinical analyses. E.S.W. performed viral burden analysis and performed immune cell processing for flow cytometry. E.S.W. and A.L.B. performed pulmonary mechanics analysis. E.S.W. and J.B.C. performed neutralization analyses J.Y. and R.H. performed RNA sequencing and analysis. H.J., Y.H., J.D.A., and T.M.R. performed hamster experiments. T.L.D. and A.C.M.B designed and validated reagents for hamster gene expression. M.M. provided anti-CCR2 antibody. J.E.C., T.M.R., A.C.M.B., and M.S.D. obtained funding and supervised research. E.S.W. and M.S.D. wrote the initial draft, with the other authors providing editorial comments.

## COMPETING FINANCIAL INTERESTS

M.S.D. is a consultant for Inbios, Vir Biotechnology, NGM Biopharmaceuticals, Carnival Corporation and on the Scientific Advisory Boards of Moderna and Immunome. The Diamond laboratory has received unrelated funding support in sponsored research agreements from Moderna, Vir Biotechnology, and Emergent BioSolutions. J.E.C. has served as a consultant for Eli Lilly and Luna Biologics, is a member of the Scientific Advisory Boards of CompuVax and Meissa Vaccines and is Founder of IDBiologics. The Crowe laboratory at Vanderbilt University Medical Center has received sponsored research agreements from AstraZeneca and IDBiologics. Vanderbilt University has applied for patents related to antibodies described in this paper. The Boon laboratory has received unrelated funding support in sponsored research agreements from AI Therapeutics, GreenLight Biosciences, AbbVie, and Nano Targeting & Therapy Biopharma.

## SUPPLEMENTAL FIGURE AND TABLE LEGENDS

**Figure S1. Protective effects of COV2-2381 or COV2-2072. Related to Figure 2.** (**A**) Anti-SARS-CoV-2 mAbs (COV2-2381 or COV2-2381 LALA) were incubated with 10^2^ focus-forming units (FFU) of SARS-CoV-2 for 1 h at 37°C followed by addition of mAb-virus mixture to Vero E6 cells. Wells containing mAb were compared to wells without MAb to determine relative infection. One experiment of two is shown. (**B**) Binding of COV2-2381 and COV2-2381 LALA to recombinant mouse FcγRI and FcγRIV as measured by ELISA (two experiments). For **C-D,** eight-week-old male K18-hACE2 transgenic mice were inoculated by the intranasal route with 10^3^ PFU of SARS-CoV-2. At 1 dpi (D+1), mice were given a single 200 μg dose of COV2-2381 or COV2-2381 LALA by intraperitoneal injection. (**C**) Weight change (mean ± SEM; n = 7-8, two experiments: two-way ANOVA with Sidak’s post-test: * *P* < 0.05, ** *P* < 0.01, **** *P* < 0.0001; comparison to the isotype control mAb-treated group). (**D**) Viral RNA levels at 8 dpi in the lung as determined by qRT-PCR (n = 8, two experiments, one-way ANOVA with Dunnett’s test; **** P < 0.0001). (**E**) Anti-SARS-CoV-2 mAbs (COV2-2072 or COV2-2072 LALA-PG) were incubated with 10^2^ FFU of SARS-CoV-2 for 1 h at 37°C followed by addition of mAb-virus mixture to Vero E6 cells. One experiment of two, with similar results, is shown. The mean of two technical replicates is shown. **F-G.** Eight-week-old male K18-hACE2 transgenic mice were inoculated by the intranasal route with 10^3^ PFU of SARS-CoV-2. At 1 (D+1) dpi, mice were given a single 200 μg dose of COV2-2072 or COV2-2072 LALA-PG by intraperitoneal injection. (**F**) Weight change (mean ± SEM; n = 6, two experiments: two-way ANOVA with Sidak’s post-test: * *P* < 0.05, ** *P* < 0.01, *** *P* < 0.001, **** *P* < 0.0001; comparison to the isotype control mAb-treated group). (**G**) Viral RNA levels at 8 dpi in the lung as determined by qRT-PCR (n = 6, two experiments: one-way ANOVA with Turkey’s post-test: ns not significant, * *P* < 0.05, *** *P* < 0.001, **** *P* < 0.0001, comparison to the isotype control mAb-treated group).

**Figure S2. Cytokine induction following SARS-CoV-2 infection. Related to Figure 3.** Cytokine levels as measured by multiplex platform in the lungs of SARS-CoV-2 infected mice at 8 dpi following isotype, COV2-2050, or COV2-2050 LALA-PG treatment at 1 dpi (D+1) or 2 dpi (D+2) (three experiments, n = 5 per group. One-way ANOVA with Dunnett’s test; * *P* < 0.05; ** *P* < 0.01; *** *P* < 0.001.) Asterisks indicate statistical significance compared to the isotype-control mAb-treated group.

**Figure S3. Flow cytometric gating strategy for BAL analysis. Related to Figure 3.** For BAL staining, cells were gated to live, single, autofluorescent-negative CD45^+^ cells to identify hematopoietic cells. Alveolar macrophages were identified as SiglecF^hi^ CD11c^hi^ cells. Neutrophils were identified as Ly6G^hi^ CD11b^hi^ cells. CD11b^-^ cells were gated further into CD4^+^ and CD8^+^ T cells. CD11b^hI^Ly6G^-^ cells were gated subsequently using CD64, CD24, and MHC-II. MHCII^hi^ CD24^hi^ were defined as CD11b^+^ DCs. MHCI^lo^ Ly6^hi^ cells were defined as monocytes.

**Figure S4. Gene ontology analysis of RNAseq data. Related to Figure 4. A-C.** RNA sequencing analysis from the lung homogenates of naive K18-hACE2 mice or mice inoculated with SARS-CoV-2 at 8 dpi. At 1 (D+1) or 2 (D+2) dpi, mice were given a single 200 μg dose of COV2-2050 or COV2-2050 LALA-PG by intraperitoneal injection. (**A-B**) Gene Ontology (GO) Enrichment Analysis of biological process terms enriched in down-regulated genes from comparisons of isotype control, COV2-2050, and COV2-2050 LALA-PG when given at D+1 (**A**) or isotype control, COV2-2050, and COV2-2050 LALA-PG when given at D+2 (**B**). Terms were ranked by the false discovery rate (q-value), and the top 20 are listed after eliminating redundant terms. (**C**) Heat maps of significantly downregulated gene sets corresponding with intact COV2-2050 treatment identified through GO analysis. Genes shown in each pathway are the union of the differentially expressed genes (DEGs) from the five comparisons (isotype control, COV2-2050 D+1, COV2-2050 D+2, COV2-2050 LALA-PG D+1, or COV2-2050 LALA-PG D+2 versus mock-infected). Columns represent samples and rows represent genes. Gene expression levels in the heat maps are z score-normalized values determined from log2cpm values.

**Figure S5. CompBio Analysis comparing COV2-2050 D+2, Isotype control, and COV2-2050 LALA-PG D+2. Related to Figure 4.** Heatmaps of selected relevant biological themes enriched in COV2-2050 D+2 versus isotype control and COV2-LALA-PG D+2. Genes shown are common in the pair of comparisons with an enrichment score of 100 or greater in either of the paired comparisons.

**Figure S6. Confirmation of cellular depletions. Related to Figure 5. A.** Representative flow cytometry plots of monocytes and neutrophils from peripheral blood at 8 dpi following intraperitoneal injection of a depleting anti-CCR2 mAb or isotype control mAb. **B.** Frequency of Ly6C^hi^ monocytes and neutrophils in blood at 8 dpi following anti-CCR2 or isotype control mAb administration in isotype control or COV2-2050-treated mice (two experiments, n = 6 per group). **C.** Representative flow cytometry plots of peripheral blood at 8 dpi following intraperitoneal injection of a depleting anti-Ly6G mAb or isotype control mAb. **D.** Frequency of Ly6C^hi^ monocytes and neutrophils in blood at 8 dpi following anti-Ly6G or isotype control mAb administration in isotype control or COV2-2050-treated mice (two experiments, n = 5-6 per group). **E**. Representative flow cytometry plots of peripheral blood at 8 dpi following intraperitoneal injection of a depleting anti-NK1.1 mAb or isotype control mAb. **D.** Frequency of NK cells in blood at 8 dpi following anti-NK1.1 or isotype control mAb administration in isotype control or COV2-2050-treated mice (two experiments, n = 4-5 per group).

**Table S1. Related to Figure 4. Gene lists associated with GO analysis.** Lists of up-regulated and down-regulated genes comparing Isotype control mAb versus COV2-2050 D+1, COV2-2050 D+1 versus COV2-2050 LALA-PG D+1, Isotype control mAb versus COV2-2050 D+2, and COV2-2050 D+2 versus COV2-2050 LALA-PG D+2 in the top pathways identified through Gene Ontology analysis. Associated q-value and fold-change values are shown.

**Table S2. Related to Figure 4. Gene lists associated with CompBio analysis.** Lists of up-regulated and down-regulated genes comparing Isotype control mAb versus COV2-2050 D+1, COV2-2050 D+1 versus COV2-2050 LALA-PG D+1, Isotype control mAb versus COV2-2050 D+2, and COV2-2050 D+2 versus COV2-2050 LALA-PG D+2 in the top pathways identified through CompBio analysis. Associated q-value and fold-change values are shown.

## STAR METHODS

### RESOURCE AVAILABLITY

#### Lead Contact

Further information and requests for resources and reagents should be directed to the Lead Contact, Michael S. Diamond.

#### Materials Availability

All requests for resources and reagents should be directed to the Lead Contact author. This includes mice, antibodies, viruses, and proteins. All reagents will be made available on request after completion of a Materials Transfer Agreement.

#### Data and code availability

All data supporting the findings of this study are available within the paper and are available from the corresponding author upon request. RNA sequencing datasets have been uploaded and are available at GSE161615.

### EXPERIMENTAL MODEL AND SUBJECT DETAILS

#### Cells and viruses

Vero E6 (CRL-1586, American Type Culture Collection (ATCC), Vero CCL81 (ATCC), and Vero-furin cells (Mukherjee et al., 2016) were cultured at 37°C in Dulbecco’s Modified Eagle medium (DMEM) supplemented with 10% fetal bovine serum (FBS), 10 mM HEPES pH 7.3, 1 mM sodium pyruvate, 1× non-essential amino acids, and 100 U/ml of penicillin-streptomycin. The 2019n-CoV/USA_WA1/2019 isolate of SARS-CoV-2 was obtained from the US Centers for Disease Control (CDC). Infectious stocks were propagated by inoculating Vero CCL81 cells and collecting supernatant upon observation of cytopathic effect; debris was removed by centrifugation and passage through a 0.22 μm filter. Supernatant was aliquoted and stored at −80°C. All work with infectious SARS-CoV-2 was performed in Institutional Biosafety Committee approved BSL3 and A-BSL3 facilities at Washington University School of Medicine using appropriate positive pressure air respirators and protective equipment.

#### Antibodies

The human antibodies studied in this paper were isolated from blood samples from two individuals in North America with previous laboratory-confirmed symptomatic SARS-CoV-2 infection that was acquired in China. The original clinical studies to obtain specimens after written informed consent were previously described (Zost et al., 2020b) and had been approved by the Institutional Review Board of Vanderbilt University Medical Center, the Institutional Review Board of the University of Washington and the Research Ethics Board of the University of Toronto.

#### Mouse experiments

Animal studies were carried out in accordance with the recommendations in the Guide for the Care and Use of Laboratory Animals of the National Institutes of Health. The protocols were approved by the Institutional Animal Care and Use Committee at the Washington University School of Medicine (assurance number A3381–01). Virus inoculations were performed under anesthesia that was induced and maintained with ketamine hydrochloride and xylazine, and all efforts were made to minimize animal suffering.

Heterozygous K18-hACE c57BL/6J mice (strain: 2B6.Cg-Tg(K18-ACE2)2Prlmn/J) were obtained from The Jackson Laboratory. Animals were housed in groups and fed standard chow diets. Eight-to nine-week-old mice of both sexes were administered 10^3^ PFU of SARS-CoV-2 by intranasal administration.

#### Hamster experiments

Seven-month-old female Syrian hamsters were purchased from Charles River Laboratories and housed in microisolator units. All hamsters were allowed free access to food and water and cared for under United States Department of Agriculture (USDA) guidelines for laboratory animals. Hamsters were administered with 5 × 10^5^ PFU of SARS-CoV-2 (2019-nCoV/USA-WA1/2020) by the intranasal route in a final volume of 100 μL. One day later, hamsters were administered by intraperitoneal injection COV2-2050, COV2-2050 LALA-PG, or isotype control (10 mg/kg). All hamsters were monitored for body weight loss until humanely euthanized at 6 dpi. All procedures were approved by the University of Georgia Institutional Animal Care and Use Committee (IACUC number A2020 04-024-Y1-A4). Virus inoculations and antibody transfers were performed under anesthesia that was induced and maintained with 5% isoflurane. All efforts were made to minimize animal suffering.

### METHOD DETAILS

#### MAb production and purification

COV2-2050, COV2-2381, COV2-2072, and COV2-3025 were isolated from a B cell from a SARS-CoV-2 convalescent patient and described previously (Zost et al., 2020a; Zost et al., 2020b). mAb 2D22 was described previously (Fibriansah et al., 2015). Sequences of mAbs that had been synthesized (Twist Bioscience) and cloned into an IgG1 monocistronic expression vector (designated as pTwist-mCis_G1) or IgG1 monocistronic expression vector containing L234A, L235A, and P329G mutations in the Fc region of the heavy chain (designated as pTwist-mCis_LALA-PG) were used for mAb secretion in mammalian cell culture. This vector contains an enhanced 2A sequence and GSG linker that allows the simultaneous expression of mAb heavy and light chain genes from a single construct upon transfection (Chng et al., 2015). mAb proteins were produced after transient transfection using the Gibco ExpiCHO Expression System (ThermoFisher Scientific) following the manufacturer’s protocol. Culture supernatants were purified using HiTrap MabSelect SuRe columns (Cytiva, formerly GE Healthcare Life Sciences) on an AKTA Pure chromatographer (GE Healthcare Life Sciences). Purified mAbs were buffer-exchanged into PBS, concentrated using Amicon Ultra-4 50-kDa centrifugal filter units (Millipore Sigma) and stored at −80 °C until use. Purified mAbs were tested routinely for endotoxin levels (found to be less than 30 EU per mg IgG). Endotoxin testing was performed using the PTS201F cartridge (Charles River), with a sensitivity range from 10 to 0.1 EU per ml, and an Endosafe Nexgen-MCS instrument (Charles River). Chinese hamster ovary cell expressed recombinant forms of COV2-2381 IgG1 or COV2-2381 IgG1 containing L234A, L235A mutations in the Fc region of the heavy chain (COV2-2381-LALA) were kindly provided by Ron Cobb, Ology Bioservices and Chris Earnhart, U.S. Joint Program Executive Office for Chemical, Biological, Radiological and Nuclear Defense.

#### Plaque forming assay

Vero-furin cells (Mukherjee et al., 2016) were seeded at a density of 2.5×10^5^ cells per well in flat-bottom 12-well tissue culture plates. The following day, medium was removed and replaced with 200 μL of 10-fold serial dilutions of the material to be titered, diluted in DMEM+2% FBS. One hours later, 1 mL of methylcellulose overlay was added. Plates were incubated for 72 h, then fixed with 4% paraformaldehyde (final concentration) in phosphate-buffered saline for 20 min. Plates were stained with 0.05% (w/v) crystal violet in 20% methanol and washed twice with distilled, deionized water.

#### Measurement of viral burden

Tissues were weighed and homogenized with zirconia beads in a MagNA Lyser instrument (Roche Life Science) in 1000 μL of DMEM media supplemented with 2% heat-inactivated FBS. Tissue homogenates were clarified by centrifugation at 10,000 rpm for 5 min and stored at −80°C. RNA was extracted using the MagMax mirVana Total RNA isolation kit (Thermo Scientific) on the Kingfisher Flex extraction robot (Thermo Scientific). RNA was reverse transcribed and amplified using the TaqMan RNA-to-CT 1-Step Kit (ThermoFisher). Reverse transcription was carried out at 48°C for 15 min followed by 2 min at 95°C. Amplification was accomplished over 50 cycles as follows: 95°C for 15 s and 60°C for 1 min. Copies of SARS-CoV-2 N gene RNA in samples were determined using a previously published assay (Case et al., 2020). Briefly, a TaqMan assay was designed to target a highly conserved region of the N gene (Forward primer: ATGCTGCAATCGTGCTACAA; Reverse primer: GACTGCCGCCTCTGCTC; Probe: /56-FAM/TCAAGGAAC/ZEN/AACATTGCCAA/3IABkFQ/). This region was included in an RNA standard to allow for copy number determination down to 10 copies per reaction. The reaction mixture contained final concentrations of primers and probe of 500 and 100 nM, respectively.

#### Cytokine and chemokine mRNA measurements

RNA was isolated from lung homogenates as described above. cDNA was synthesized from DNAse-treated RNA using the High-Capacity cDNA Reverse Transcription kit (Thermo Scientific) with the addition of RNase inhibitor following the manufacturer’s protocol. Mouse cytokine and chemokine expression was determined using TaqMan Fast Universal PCR master mix (Thermo Scientific) with commercial primers/probe sets specific for *IFN-g* (IDT: Mm.PT.58.41769240), *IL-6* (Mm.PT.58.10005566), *CXCL10* (Mm.PT.58.43575827), *CCL2* (Mm.PT.58.42151692), and results were normalized to *GAPDH* (Mm.PT.39a.1) levels. Fold change was determined using the 2^-ΔΔCt^ method comparing treated mice to naïve controls. Hamster cytokine and chemokine expression was determined using TaqMan Fast Universal PCR master mix (Thermo Scientific) with primers/probe sets specific for *Rpl18* (For: GTTTATGAGTCGCACTAACCG, Rev: TGTTCTCTCGGCCAGGAA, Probe: YAK-TCTGTCCCTGTCCCGGATGATC-BBQ), *Cxcl10* (For: GCCATTCATCCACAGTTGACA, Rev: CATGGTGCTGACAGTGGAGTCT, Probe: 6FAM-CGTCCCGAGCCAGCCAACGA-BBQ), *Ccl2* (For: CTCACCTGCTGCTACTCATTC, Rev: CTCTCTCTTGAGCTTGGTGATG, Probe: 6FAM-CAGCAGCAAGTGTCCCAAAGAAGC-BBQ), *Ccl3* (For: CTCACCTGCTGCTACTCATTC, Rev: CTCTCTCTTGAGCTTGGTGATG, Probe: 6FAM-CAGCAGCAAGTGTCCCAAAGAAGC-BBQ), *Ccl5* (For: TGCTTTGACTACCTCTCCTTTAC, Rev: GGTTCCTTCGGGTGACAAA, Probe: 6FAM-TGCCTCGTGTTCACATCAAGGAGT-BBQ), *Ifit3* (For: CTGATACCAACTGAGACTCCTG, Rev: CTTCTGTCCTTCCTCGGATTAG, Probe: 6FAM-ACCGTACAGTCCACACCCAACTTT-BBQ). Fold change was determined using the 2^-ΔΔCt^ method comparing treated hamster to naïve controls.

#### Cytokine and chemokine protein measurements

Lung homogenates were incubated with Triton-X-100 (1% final concentration) for 1 h at room temperature to inactivate SARS-CoV-2. Homogenates then were analyzed for cytokines and chemokines by Eve Technologies Corporation (Calgary, AB, Canada) using their Mouse Cytokine Array / Chemokine Array 44-Plex (MD44) platform.

#### Lung histology

Animals were euthanized before harvest and fixation of tissues. The left lung was first tied off at the left main bronchus and collected for viral RNA analysis. The right lung then was inflated with ~1.2 mL of 10% neutral buffered formalin using a 3-mL syringe and catheter inserted into the trachea. Tissues were embedded in paraffin, and sections were stained with hematoxylin and eosin. Tissue sections were visualized using a Nikon Eclipse microscope equipped with an Olympus DP71 camera, a Leica DM6B microscope equipped with a Leica DFC7000T camera, or an Olympus BX51 microscope with attached camera.

#### Flow cytometry analysis of immune cell infiltrates

For analysis of BAL fluid, mice were sacrificed by ketamine overdose, followed by cannulation of the trachea with a 19-G canula. BAL was performed with three washes of 0.8 ml of sterile PBS. BAL fluid was centrifuged, and single cell suspensions were generated for staining. Single cell suspensions of BAL were preincubated with Fc Block antibody (BD PharMingen) in PBS + 2% heat-inactivated FBS for 10 min at room temperature before staining. Cells were incubated with antibodies against the following markers: AF700 anti-CD45 (clone 30 F-11), APC-Cy7 anti-CD11c (clone N418), PE anti-Siglec F (clone E50-2440; BD), PE-Cy7 anti-Ly6G (clone 1A8), BV605 anti-Ly6C (clone HK1.4; BioLegend), BV 711 anti-CD11b (clone M1/70), APC anti-CD103 (clone 2E7; eBioscience), PB anti-CD3 (clone 17A2), PE-Cy7, APC anti-CD4 (clone RM4-5), PE-Cy7 anti-CD8 (clone53-6.7), anti-NK1.1 (clone PK136), and BV605 anti-TCR γ/δ (clone GL3). All antibodies were used at a dilution of 1:200. Cells were stained for 20 min at 4°C, washed, fixed and permeabilized for intracellular staining with Foxp3/Transcription Factor Staining Buffer Set (eBioscience) according to manufacturer’s instructions. Cells were incubated overnight at 4°C with PE-Cy5 anti-Foxp3 (clone FJK-16s), washed, re-fixed with 4% PFA (EMS) for 20 min and resuspended in permeabilization buffer. Absolute cell counts were determined using Trucount beads (BD). Flow cytometric data were acquired on a cytometer (BD-X20; BD Biosciences) and analyzed using FlowJo software (Tree Star).

#### Antibody depletion of immune cell subsets

For neutrophil depletion, anti-Ly6G (BioXCell; clone 1A8; 250 μg) or an isotype control (BioXCell; clone 2A3; 250 μg) was administered to mice by intraperitoneal injection one day before infection and at D+1, D+3, and D+6 relative to SARS-CoV-2 infection. For monocyte depletion, anti-CCR2 (clone MC-21; 50 μg) (Mack et al., 2001) or an isotype control mAb (BioXCell; clone LTF-2; 50 μg) was administered to mice by intraperitoneal injection one day before infection and at D+1, D+3, D+5, and D+7 relative to SARS-CoV-2 infection. For NK cell depletion, anti-NK1.1 (clone PK136; 200 μg) or an isotype control mAb (BioXCell; clone C1.18.4; 200 μg) was administered to mice by intraperitoneal injection one day before infection and at D+1, D+3, D+5, and D+7 relative to SARS-CoV-2 infection.

For analysis of immune cell depletion, peripheral blood was collected on the day of harvest. Erythrocytes were lysed with ACK lysis buffer (Gibco) and resuspended in RPMI supplemented with 10% FBS. For anti-Ly6G or anti-CCR2 depletion, single-cell suspensions were preincubated with Fc Block antibody (BD PharMingen) in PBS + 2% heat-inactivated FBS for 10 min at room temperature and then stained with antibodies against CD45 BUV395, CD11b PE/Dazzle 594, Ly6C Pacific Blue, Ly6B FITC, Ly6G BV650, and fixable viability dye (eFluor 506). For NK cell depletions, single cell suspensions were blocked for FcγR binding and stained with antibodies against CD45 BUV395, CD19 BV711, CD3 BV711, NK1.1 FITC, and Nkp56 Pacific Blue.

#### Respiratory mechanics

Mice were anesthetized with ketamine/xylazine (100 mg/kg and 10 mg/kg, i.p., respectively). The trachea was isolated by dissection of the neck area and cannulated using an 18-gauge blunt metal cannula (typical resistance of 0.18 cmH_2_O.s/mL), which was secured in place with a nylon suture. The mouse then was connected to the flexiVent computer-controlled piston ventilator (SCIREQ Inc.) via the cannula, which was attached to the FX adaptor Y-tubing. Mechanical ventilation was initiated, and mice were given an additional 100 mg/kg of ketamine and 0.1 mg/mouse of the paralytic pancuronium bromide by intraperitoneal route to prevent breathing efforts against the ventilator and during measurements. Mice were ventilated using default settings for mice, which consisted in a positive end expiratory pressure at 3 cm H_2_O, a 10 mL/kg tidal volume (Vt), a respiratory rate at 150 breaths per minute (bpm), and a fraction of inspired oxygen (FiO_2_) of 0.21 (*i.e*., room air). Respiratory mechanics were assessed using the forced oscillation technique, as previously described (McGovern et al., 2013), using the latest version of the flexiVent operating software (flexiWare v8.1.3). Pressure-volume loops and measurements of inspiratory capacity also were done.

#### Neutralization assay

Serial dilutions of mAbs were incubated with 10^2^ focus-forming units (FFU) of SARS-CoV-2 for 1 h at 37°C. mAb-virus complexes were added to Vero E6 cell monolayers in 96-well plates and incubated at 37°C for 1 h. Subsequently, cells were overlaid with 1% (w/v) methylcellulose in MEM supplemented with 2% FBS. Plates were harvested 30 h later by removing overlays and fixed with 4% PFA in PBS for 20 min at room temperature. Plates were washed and sequentially incubated with 1 μg/mL of a recombinant IgG based on the sequence of CR3022 (Yuan et al., 2020) anti-S antibody and HRP-conjugated goat anti-human IgG in PBS supplemented with 0.1% saponin and 0.1% bovine serum albumin. SARS-CoV-2-infected cell foci were visualized using TrueBlue peroxidase substrate (KPL) and quantitated on an ImmunoSpot microanalyzer (Cellular Technologies).

#### RNA sequencing

cDNA libraries were constructed starting with 10 ng of total RNA from lung tissues of each sample that was extracted using the MagMax Mirvana kit per manufacturer’s instructions. cDNA was generated using the Seqplex kit (Sigma-Aldrich, St. Louis, MO) with amplification of 20 cycles. Library construction was performed using 100 ng of cDNA undergoing end repair, A-tailing, ligation of universal TruSeq adapters, and 8 cycles of amplification to incorporate unique dual index sequences. Libraries were sequenced on the NovaSeq 6000 (Illumina, San Diego, CA) targeting 40 million read pairs and extending 150 cycles with paired end reads. RNA-seq reads were aligned to the mouse Ensembl GRCh38.76 primary assembly and SARS-CoV-2 NCBI NC_045512 Wuhan-Hu-1 genome with STAR program (version 2.5.1a) (Dobin et al., 2013). Gene counts were derived from the number of uniquely aligned unambiguous reads by Subread:featureCount (version 1.4.6-p5) (Liao et al., 2014). The ribosomal fraction, known junction saturation, and read distribution over known gene models were quantified with RSeQC (version 2.6.2) (Liao et al., 2014). All gene counts were preprocessed with the R package EdgeR (Robinson et al., 2010) to adjust samples for differences in library size using the trimmed mean of M values (TMM) normalization procedure. Viral and ribosomal genes and genes not expressed in at least five samples (the smallest group size) at a level greater than or equal to 1 count per million reads were excluded, resulting 12,157 unique genes in further analysis. The R package limma (Ritchie et al., 2015) with voomWithQualityWeights function (Liu et al., 2015) was utilized to calculate the weighted likelihoods for all samples, based on the observed mean-variance relationship of every gene and sample. Differentially expressed genes were defined as those with at least 2-fold difference between two individual groups at the Benjamini-Hochberg false-discovery rate (FDR) (https://www.jstor.org/stable/2346101?seq=1) adjusted *P* value (i.e. q-value < 0.05).

### QUANTIFICATION AND STATISTICAL ANALYSIS

Statistical significance was assigned when *P* values were < 0.05 using Prism version 8 (GraphPad). Tests, number of animals (n), median values, and statistical comparison groups are indicated in the Figure legends. Analysis of weight change was determined by two-way ANOVA. Changes in functional parameters or immune parameters were compared to isotype-treated animals and were analyzed by one-way ANOVA or one-way ANOVA with Dunnett’s test.

