## Supplementary figures and images for "Human neutralizing antibodies against SARS-CoV-2 require intact Fc effector functions and monocytes for optimal therapeutic protection"

### Supplemental Figures 1 to 6

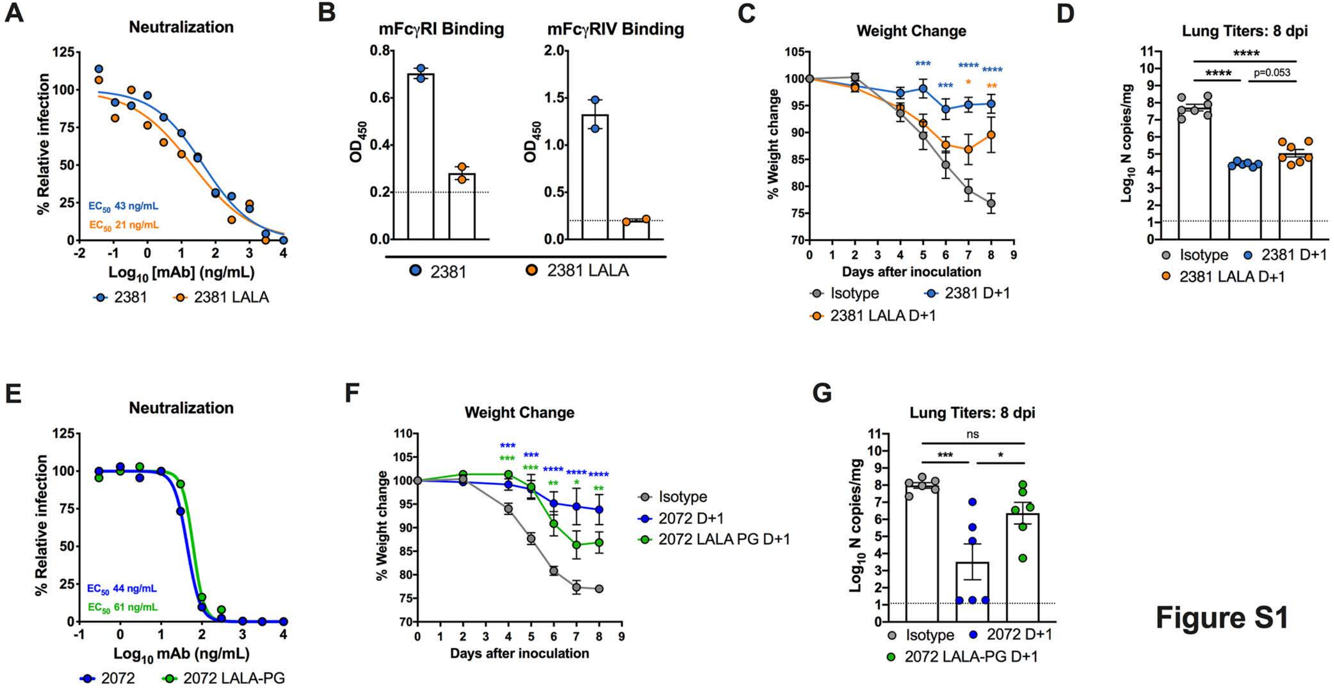

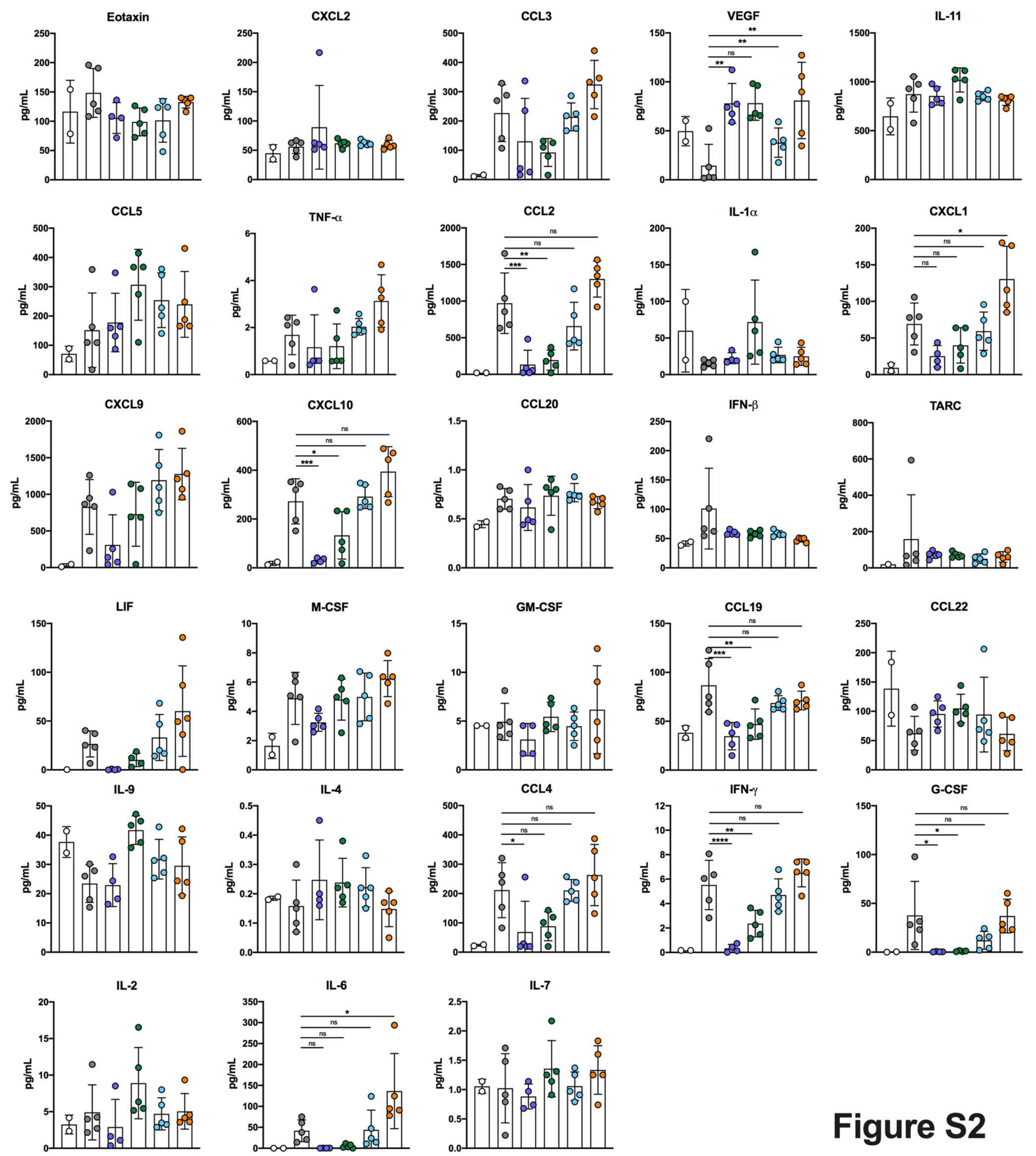

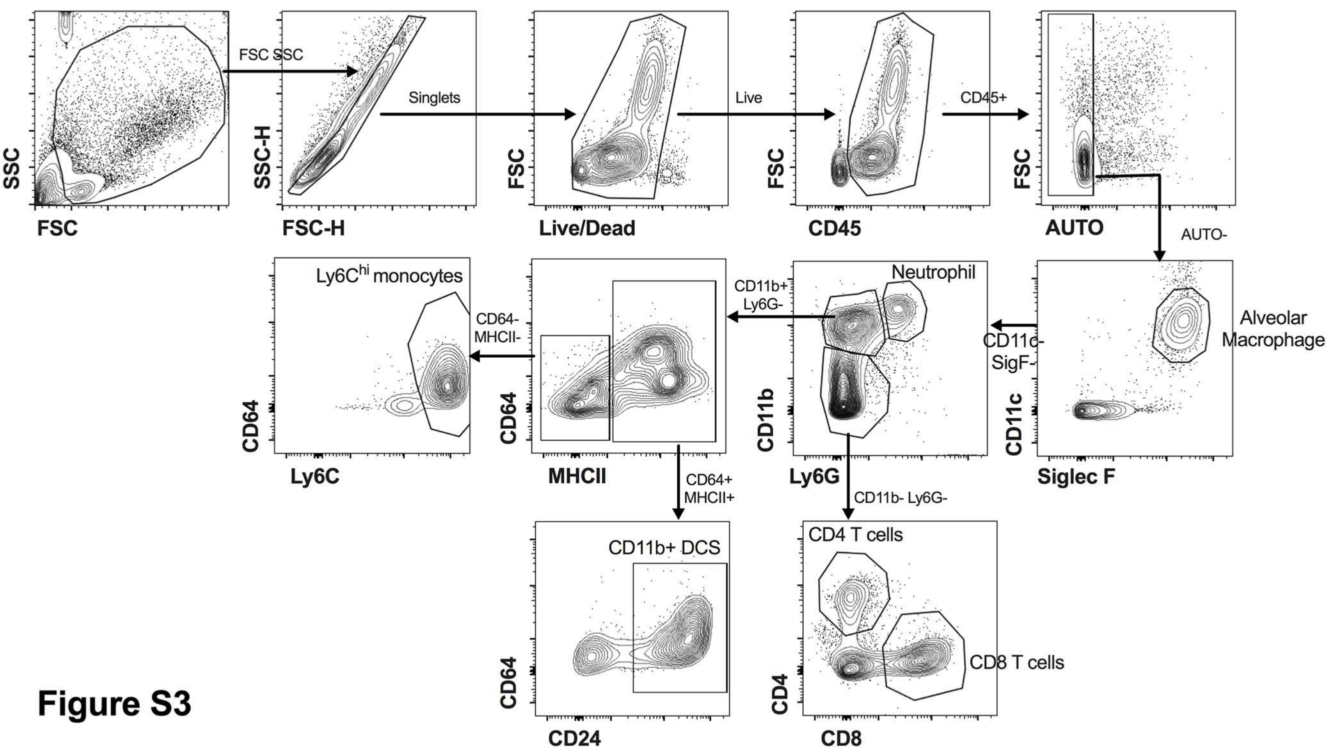

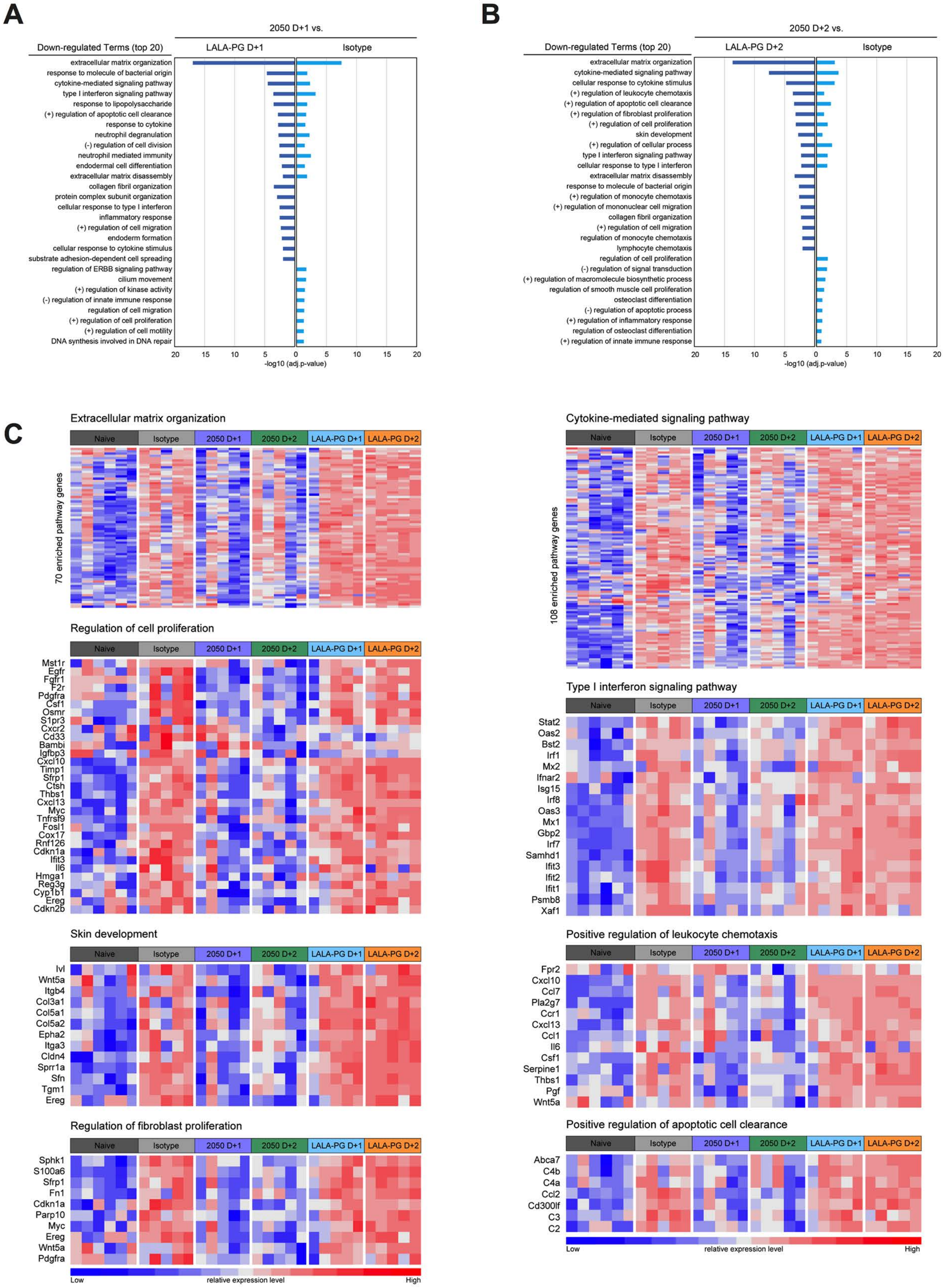

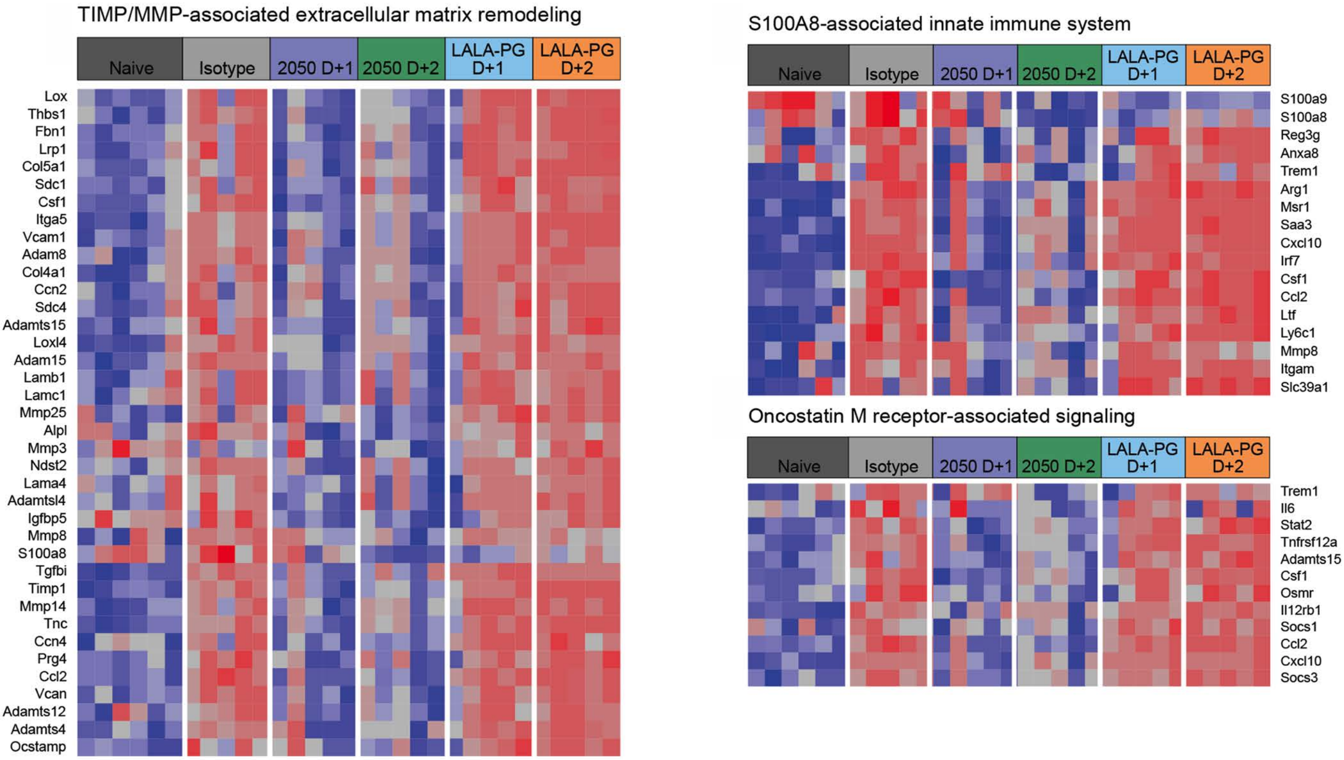

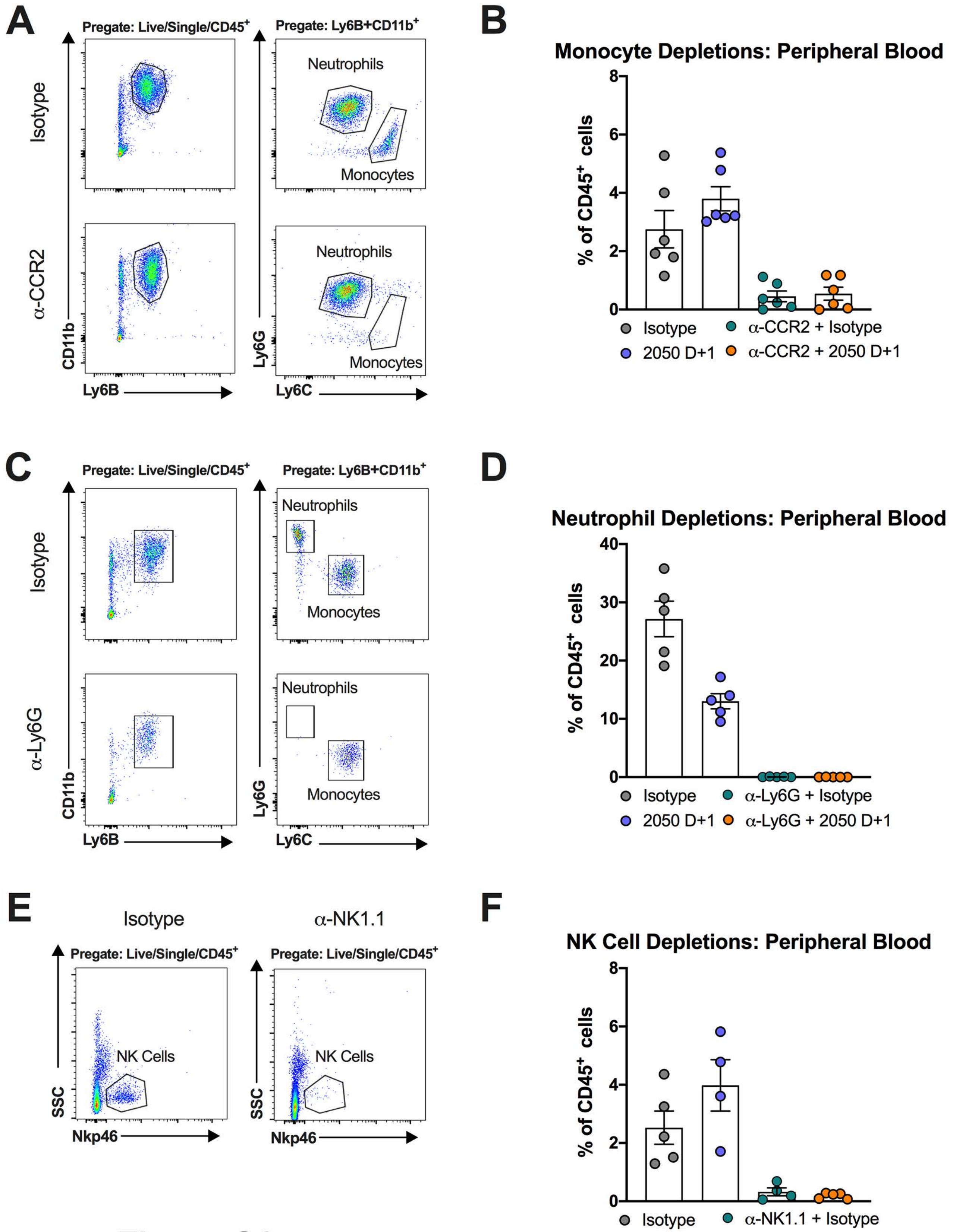
